# Spatial Tumour Tissue Distribution Dynamics of a Bicycle Peptide-MMAE Toxin Conjugate Profiled by Targeted Mass Spectrometry

**DOI:** 10.64898/2026.10.13.758755

**Authors:** Maria-Konstantina Ioannidi, Simon Davis, Tianen He, Svenja Hester, Mike Rigby, Finn Holding, Phil Jeffrey, Michael Skynner, Iolanda Vendrell, Roman Fischer, Benedikt M. Kessler

## Abstract

A highly specific-NECTIN-4 binding bicyclic peptide covalently linked to the cytotoxin monomethyl auristatin E (MMAE) via a valine-citrulline cleavable linker (zelenectide pevedotin, BT8009) is a promising modality to treat cancer patients with solid tumours. Although the pharmacokinetics and biodistribution of zelenectide pevedotin have been well characterised, a more precise view of tumour delivery and intra-tumoural distribution of its released payload is lacking. To address this, we have developed an approach to profile the spatial tumour and healthy tissue distribution of this anti-cancer agent using laser capture microdissection coupled to targeted liquid chromatography parallel reaction monitoring (LCM-LC-PRM) mass spectrometry. This LCM-assisted spatial pharmacology workflow enables the precise isolation of specific tissue regions, followed by high-resolution analysis of MMAE distribution at sub-tissue-specific levels. By identifying and optimising MS/MS-specific fragmentation ions for accurate and sensitive detection and quantitation of MMAE released from zelenectide pevedotin within tumour tissues, we observed progressive accumulation and sustained retention of MMAE in mouse tumour tissues over a period of 24 h. Conversely, in plasma, liver and muscle tissues, MMAE levels generally declined following early exposure, although inter-animal variability was observed. LCM-based intra-tumour spatial profiling revealed heterogeneous MMAE distribution between tumour cell-rich and stroma-rich regions, with distinct temporal patterns across compartments. Together, our spatial targeted MS method provided valuable insights into zelenectide pevedotin’s payload distribution dynamics *in vivo*, thereby enhancing our understanding of its therapeutic potential and guiding the development of next-generation Bicycle Drug Conjugate (BDC).

## INTRODUCTION

Targeted delivery of cytotoxic agents to tumours via antibody-drug conjugates (ADCs) has emerged as a promising cancer treatment strategy[1]. ADCs leverage the traits of high affinity and specificity of antibodies towards cognate tumour antigens to deliver toxins, cytotoxic or cytostatic agents to cancer cells. However, their clinical success, particularly in solid tumours, has been limited. Despite encouraging preclinical data obtained over two decades since the approval of the first ADC, *gemtuzumab ozogamicin*, only a small number of ADCs have gained FDA approval for the treatment of solid tumours[2].

A key challenge limiting ADC efficacy in solid tumours is poor tumour penetration, primarily due to the large molecular size of antibodies (∼150 kDa), which constrains their pharmacokinetics and biodistribution profile[3]. These macromolecules struggle to traverse the endothelial barrier, diffuse through the tumour’s dense extracellular matrix and overcome elevated interstitial fluid pressure. As a result, only a small fraction of the administered antibody dose (typically <0.1%) reaches and accumulates within solid tumours[4]. Additionally, high antigen density in perivascular tumour regions can create a “binding site barrier”, where ADCs bind near blood vessels, preventing deeper tissue penetration[5]. This suggests that proximal tumour cells may sequester excess toxin, thereby limiting its penetration into deeper tumour regions and reducing overall treatment efficacy. Strategies such as increasing the drug-to-antibody ratio (DAR) or total dose may improve penetration, but often at the cost of increased off-target toxicity[3]. These limitations underscore the need for delivery systems that enhance tumour distribution while minimising systemic exposure.

Targeting selective tumour antigens for effective drug delivery is also a key factor. For instance, NECTIN-4, a cell adhesion molecule overexpressed in various cancers and associated with poor prognosis, has emerged as a valuable target[6,7]. ADCs directed against NECTIN-4, such as *enfortumab vedotin* (EV), deliver the microtubule-disrupting toxin monomethyl auristatin E (MMAE) via a cleavable valine-citrulline linker. EV has demonstrated efficacy in urothelial carcinoma and received FDA approval in 2021 for patients with locally advanced or metastatic urothelial cancer, following prior treatments[8,9].

To address the size and penetration limitations of ADCs, peptide-drug conjugates have been developed as alternatives[10], where the peptide part mimics the complementarity-determining region (CDR) of the antibody[11]. Bicyclic peptides (∼4-4.5 kDa) can be chemically synthesised linked to cytotoxic agents through cleavable linkers. These small molecules with inherent chemical tunability offer efficient tumour penetration, rapid distribution into extravascular space, and moderate systemic clearance, primarily via renal elimination. Bicycle drug conjugates (BDCs) generally have a 1:1 peptide-to-toxin ratio, reducing off-target exposure and allowing high drug concentrations to circumvent the binding site barrier[12]. Furthermore, unlike ADCs, BDCs do not depend entirely on internalisation for activity and may enable extracellular toxin release within the tumour microenvironment (TME), facilitating bystander killing of nearby stromal and cancer cells[12].

Zelenectide pevedotin (formerly known as BT8009) is a recently developed BDC targeting NECTIN-4, consisting of a NECTIN-4-binding peptide linked to MMAE via a sarcosine decamer spacer. The spacer reduces steric hindrance and allows for efficient enzymatic cleavage by tumour-associated proteases such as cathepsin B[13]. Importantly, zelenectide pevedotin does not require internalisation to release MMAE, potentially avoiding common ADC resistance mechanisms involving altered endocytosis or lysosomal processing[14]. In preclinical models, zelenectide pevedotin demonstrates strong tumour regression, highlighting its potential for improved efficacy and safety compared to traditional ADCs, such as EV[15]. Tumour penetration of MMAE is significant and is retained long after both parent BDC and toxin are cleared from systemic circulation. However, the degree and spatial distribution of tumour penetration remain poorly defined.

Despite the rapid evolution of targeted therapies, the spatial pharmacology of toxin distribution within solid tumours remains largely unexplored, particularly for more advanced modalities such as bicyclic peptide-drug conjugates.

Current advanced analytics specialising in localisation and imaging[16], such as spatial proteomics[17] and spatial pharmacology[18] may offer opportunities to address this in a systematic fashion. Laser capture microdissection (LCM)[19,20] has been used to determine sub-tissue tumour proteomic profiles[21] and drug abundance[22–24].

In this study, we describe the spatial profiling of the BDC zelenectide pevedotin and its released cargo MMAE in tumour tissues using an optimised workflow that combines laser capture microdissection (LCM) with liquid chromatography parallel reaction monitoring (PRM) mass spectrometry (LCM-LC-PRM). First, distinct fragmentation profiles of zelenectide pevedotin and MMAE were established and acquisition parameters optimised for sensitive detection. Second, bulk tissue analyses from aliquoted tumour and non-tumour tissues from mice exposed to zelenectide pevedotin were performed. Third, we applied LCM to isolate stroma and corresponding matched tumour regions, enabling the precise quantification of drug distribution across spatially defined regions in xenograft models. Application of this LCM-assisted spatial pharmacology approach to tumour tissue revealed heterogeneous accumulation of MMAE within tumour-cell-rich regions compared to surrounding stroma, providing high-resolution spatial characterisation of intra-tumour toxin distribution of a BDC *in vivo*.

## RESULTS

### Establishing a mass spectrometry detection method for zelenectide pevedotin

Zelenectide pevedotin is a complex molecule comprised of a hydrophobic MMAE toxin, a cleavable linker (Glut-Val-Cit-p-aminobenzyl carbamate (PABC)), a Sar₁₀-β-alanine spacer, and a bicyclic peptide scaffold (**Figures 1A and S1A**). These structural features collectively determine its fragmentation behaviour, chromatographic properties, and ionisation efficiency. Previous mass spectrometry (MS) detection methods for zelenectide pevedotin were based on intact mass measurements [12,13,25]. To improve molecular resolution and enable quantitative analysis in cellular and tissue samples, we optimised data-dependent acquisition (DDA)-MS/MS experiments to characterise the fragmentation profile of zelenectide pevedotin and identify diagnostic product ions suitable for subsequent targeted quantification.

**Figure 1.**
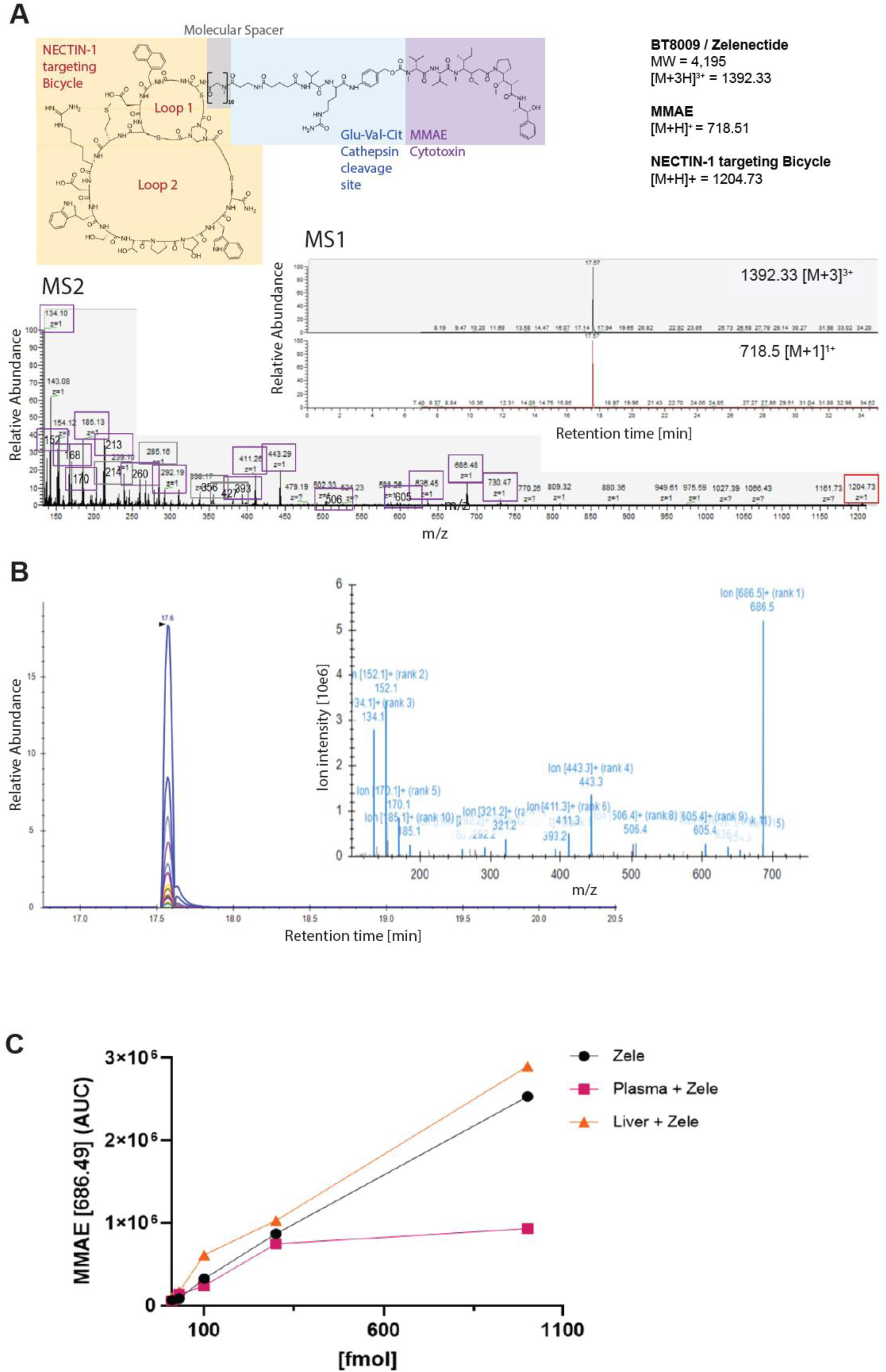
Optimising a targeted MS method for sensitive detection of zelenectide pevedotin in tissue material. **(A)** LC**-**MS/MS spectrum of BT8009 (’zelenectide pevedotin’). The majority of observed fragment masses are derived from the linker and linear peptide parts (mapped to structure, right panel). Inserts show LC-MS chromatograms of zelenectide pevedotin (top, *m/z* 1391.3 ([M + 3H]^3^⁺)) and a fragment assigned to MMAE (bottom, *m/z* 718.5 ([M + H]⁺)). **(B)** Parallel reaction monitoring (PRM) method tracking 16 MMAE characteristic fragment ions. **(C)** Calibration curves of zelenectide pevedotin standards in plasma (magenta) and liver (orange) matrices compared with solvent-based standards (black), constructed using the area under the curve (AUC) of the *m/z* 686.5 transition to evaluate matrix effects and analytical linearity. Linear regression demonstrated strong linearity for solvent standards (*R²* = 0.998) and liver matrix samples (*R²* = 0.991), whereas plasma samples exhibited reduced linearity (*R²* = 0.790), consistent with pronounced matrix effects. Calibration curves are shown on a linear scale to facilitate comparison of PRM responses across the concentration range (10-1000 nM).

For DDA MS/MS analysis, the inclusion list comprised precursor ions at *m/z* 1043.7 ([M + 4H]^4^⁺), 1391.3 ([M + 3H]^3^⁺), and 2087.5 ([M + 2H]^2^⁺), which were isolated and subjected to Higher-energy C-trap Dissociation (HCD) to generate diagnostic fragment ions for spectral characterisation. A total of 23 fragment ions were detected from the standard zelenectide pevedotin drug, of which 16 were assigned to the MMAE portion of the molecule (**Figure S1Β**). These fragments were predominantly generated by the MMAE moiety and its linker, whereas the bicyclic peptide portion remained largely intact (**Figure 1A**), underscoring its structural stability.

The identified fragment ions were assigned to defined regions of the MMAE core structure and linker system (**Figure 1, S1**), formed the basis for subsequent targeted mass spectrometry experiments utilising parallel reaction monitoring (PRM; **Figure 1B**). Specifically, low-mass fragment ions at *m/z* 134.1 and 152.1 were derived from the C-terminal aromatic region of MMAE, with the latter corresponding to norephedrine substructure, and the former resulting from the loss of a water molecule. Fragment ions at *m/z* 170.1 and 185.1 are proposed to arise from the dolaproline and dolaisoleucine units of the MMAE moiety, respectively, reflecting early cleavage events within the modified peptide chain. The *m/z* 321.2 ion is tentatively assigned to cleavage near the dolaproline ring of the MMAE moiety. These ions likely represent early-stage breakdown products near the MMAE conjugation point. In contrast, the ions observed at *m/z* 393.2, 411.2, and 443.3 correspond to intermediate-sized fragments of the MMAE structure, indicative of incremental losses along the C-terminal region of the molecule. The higher mass fragment *m/z* 506.4 is consistent with the MMAE toxin lacking the terminal carbamate group, whereas *m/z* 553.4 likely corresponds to the toxin prior to methanol elimination. Additional ions at *m/z* 605.4, 636.4, and 654.5 are tentatively assigned to MMAE-derived species exhibiting neutral losses such as water or ammonia.

The most prominent high-mass product ion was detected at *m/z* 686.5, corresponding to [M + H - CH₃OH]⁺, indicating methanol elimination from the terminal N-methyl carbamate group of the protonated intact MMAE ion ([M + H]⁺ at *m/z* 718.5). This methanol elimination is consistent with known behaviour of carbamate-containing structures and serves as the key diagnostic quantifier ion for MMAE detection. Together, these structurally annotated ions (**Figure S1**) provide a robust spectral fingerprint of MMAE within zelenectide pevedotin. Additionally, fragmentation of the Sar₁₀-β-alanine molecular spacer region produced a distinct series of five ions at *m/z* 214.1, 285.2, 356.2, 427.2, and 498.3, displaying a consistent 71 Da mass shift, reflecting sequential cleavage of the sarcosine-β-alanine repeat units (**Figure 1A**). An additional fragment at *m/z* 503.2 is tentatively assigned to the Glu-Val-Cit-PABC linker region, whereas a unique fragment at *m/z* 1204.8 likely originates from the bicyclic peptide top loop, suggesting partial preservation of this region during fragmentation.

In summary, this high-resolution mass fragmentation analysis enabled the confident assignment of zelenectide pevedotin-related ions across the MMAE, linker, and spacer regions while suggesting limited fragmentation of the bicyclic peptide scaffold. This detailed fragmentation fingerprint served as the analytical foundation for all subsequent targeted MS-based quantification and tissue localisation experiments.

### Sensitivity assessment and matrix evaluation to optimise zelenectide pevedotin detection in complex biological samples

Initial assay sensitivity for zelenectide pevedotin was assessed using pure standard solutions. A six-point calibration series spanning nominal concentrations of 1, 10, 30, 100, 300, and 1000 nM was analysed by nLC-PRM on a Q Exactive Orbitrap mass spectrometer using a 35-minute chromatographic gradient. Given the fixed injection volume of 1 μL, these standards corresponded to 1-1000 fmol on-column. Among the 16 monitored fragment ions, the transition at *m/z* 686.5 consistently produced the highest and most reproducible signal and was therefore selected as the quantifier ion throughout the study.

Linear regression across the calibration series demonstrated excellent linearity (*R²* = 0.998), with a slope (S) of 2506 and an intercept of 49422. Limits of detection (LOD) and quantitation (LOQ) were calculated according to ICH Q2(R2) guidelines using the regression standard error (*σ*), yielding an LOD of 4.76 fmol on-column and an LOQ of 14.41 fmol on-column. The lowest calibration standard (1 fmol on-column; AUC: 21,380) produced a clearly detectable signal above the LOD; however, as it fell below the defined LOQ threshold (14.41 fmol), a five-point quantitative range (10-1000 fmol on-column) was implemented for strict quantitative conversion in biological matrices. No interfering peaks were detected in blank solvent or drug-free tissue matrices, confirming the analytical specificity of the assay.

To evaluate matrix effects, zelenectide pevedotin was spiked into enzymatically digested liver homogenates and plasma obtained from untreated mice. Matching calibration series (10-1000 nM) were analysed under identical LC-PRM conditions. Comparison of solvent- and matrix-based calibration curves revealed clear matrix-dependent differences in signal response (**Figure 1C**). Liver homogenates produced consistently higher signal responses than solvent standards (*R²* = 0.991), suggesting matrix-associated signal enhancement. In contrast, plasma samples produced lower responses across the calibration range, consistent with ion suppression effects. Furthermore, the highest plasma calibration standard (1000 nM) showed a reduced incremental response relative to the 300 nM standard (934822 *vs*. 748041), suggesting a deviation from linear detector response at the highest concentration, which contributed to reduced linearity (*R²* = 0.790). These observations demonstrate that biological matrices differentially influence LC-PRM signal intensity and should therefore be considered during interpretation of subsequent quantitative *in vivo* measurements.

This calibration strategy, incorporating both solvent-based and matrix-spiked standards, facilitated interpretation of matrix-dependent signal variation during subsequent quantitative analyses. Collectively, these experiments established the analytical performance of the LC-PRM assay and defined the calibration framework used for subsequent *in vivo* pharmacokinetic analyses.

Although intact zelenectide pevedotin was readily detectable in solvent standards and matrix-spiked samples, it was only detected at very low abundance in tumour, liver, muscle and plasma samples collected from treated animals and could not be reliably quantified under the applied analytical conditions. This observation is consistent with extensive *in vivo* processing of the parent compound and/or rapid release of the MMAE payload. These findings are consistent with the expected pharmacology of cleavable bicyclic peptide-drug conjugates and support the conclusion that released MMAE represents the predominant analytically robust species *in vivo*. Accordingly, all subsequent pharmacokinetic and spatial distribution analyses focused on MMAE as a surrogate marker of zelenectide pevedotin exposure.

### *In vivo* pharmacokinetics and tissue distribution of zelenectide pevedotin (MMAE)

Following intravenous administration of zelenectide pevedotin, released MMAE was quantified in tumour, plasma, liver, and muscle samples collected from 12 female BALB/c nude mice (6-8 weeks old). Tissue and plasma samples were obtained from the same individual animals at each post-dose timepoint, enabling paired comparisons of MMAE distribution across biological compartments. Samples were collected at 1, 6, 12, and 24 h following intravenous administration of zelenectide pevedotin, together with untreated controls. Data processing was performed using Skyline software, and quantification was based on the dominant PRM fragment ion (*m/z* 686.5), selected following optimisation of the fragmentation profile of intact zelenectide pevedotin.

No drug-related fragment ions were detected in untreated control tissues, supporting the analytical specificity of the LC-PRM method (**Figure S2**). Representative chromatograms obtained from tumour tissue (**Figure 2**) and additional tissues (**Figures S3-S5**) demonstrate clear, well-resolved chromatographic peaks with consistent retention times and reproducible fragment ion profiles across biological replicates and sampling timepoints. Chromatographic co-elution of multiple qualifier ions with the quantifier transition further supported confident analyte identification throughout the study. The presence of clearly resolved peaks with baseline separation and consistent fragment ion alignment across time points reflects accurate spectral resolution and robust mass spectrometric performance. Minor variability in retention time was within the expected range for nano-flow LC conditions (<1%), further supporting the method’s precision and reproducibility under the applied analytical conditions.

**Figure 2.**
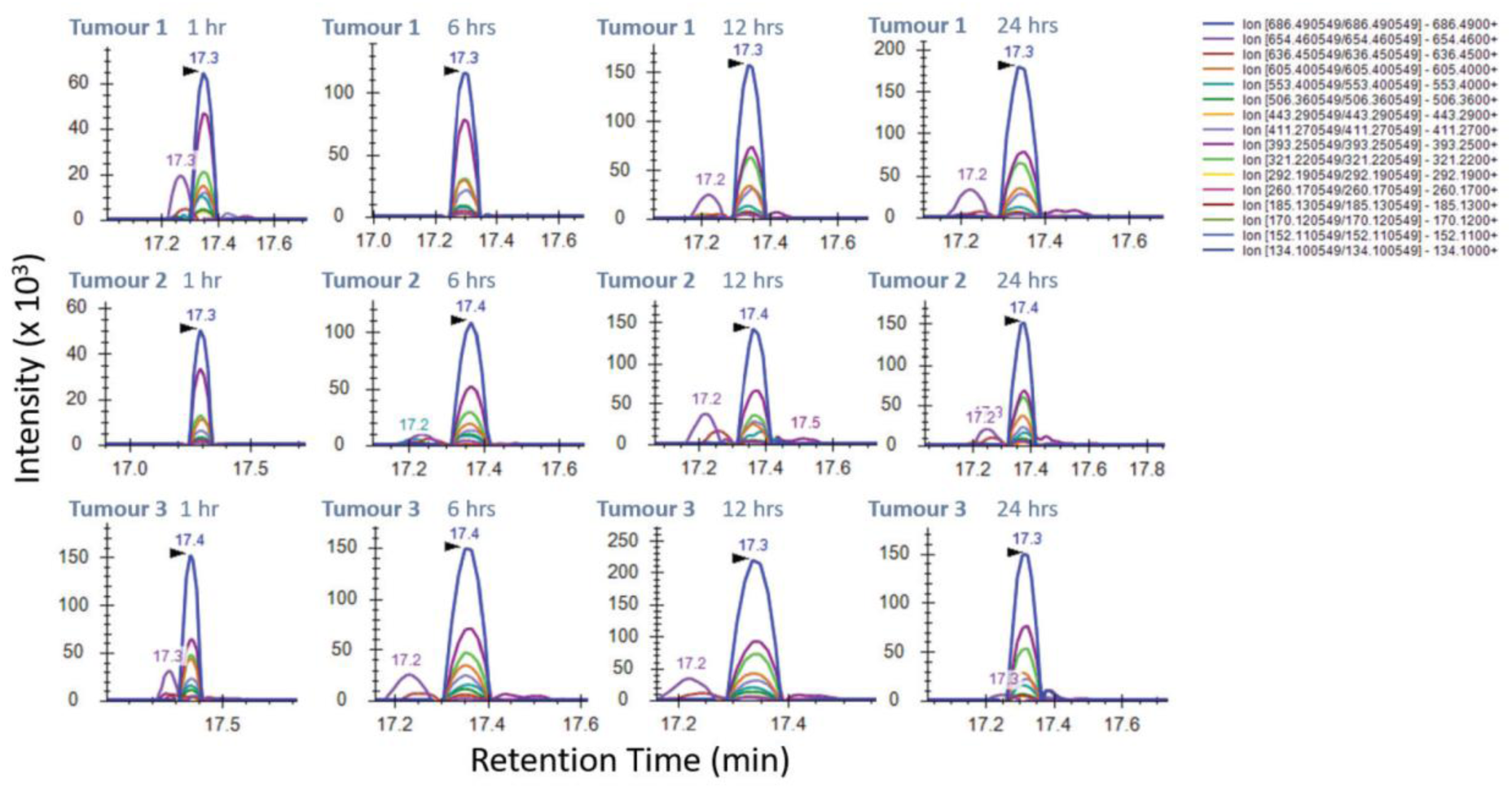
Representative LC-PRM chromatograms of MMAE detected in tumour tissue. Chromatograms show MMAE fragmentation profiles at 1, 6, 12, and 24 h post-administration of zelenectide pevedotin. The dominant quantifier fragment ion (*m/z* 686.5) and multiple qualifier ions demonstrate reproducible retention times and chromatographic alignment across biological replicates. Background signals observed in control tissue are shown in Figure S2. Chromatograms display retention time (x-axis) versus signal intensity (y-axis) for 16 fragment ions derived from the precursor ion (*m/z* 718.5). Each fragment ion is represented by a different colour, and peak areas reflect their relative concentrations, ranked from most to least abundant. Labels indicate tissue type, biological replicate, and sampling time point (e.g., ‘Tumour_1_1h’). Quantitative pharmacokinetic profiles derived from the quantifier transition are presented in Figure 3.

These chromatograms provide qualitative confirmation of MMAE detection and demonstrate consistent analytical performance across all analysed tissues. Quantitative pharmacokinetic profiles derived from the quantifier transition are presented in **Figure 3**.

**Figure 3.**
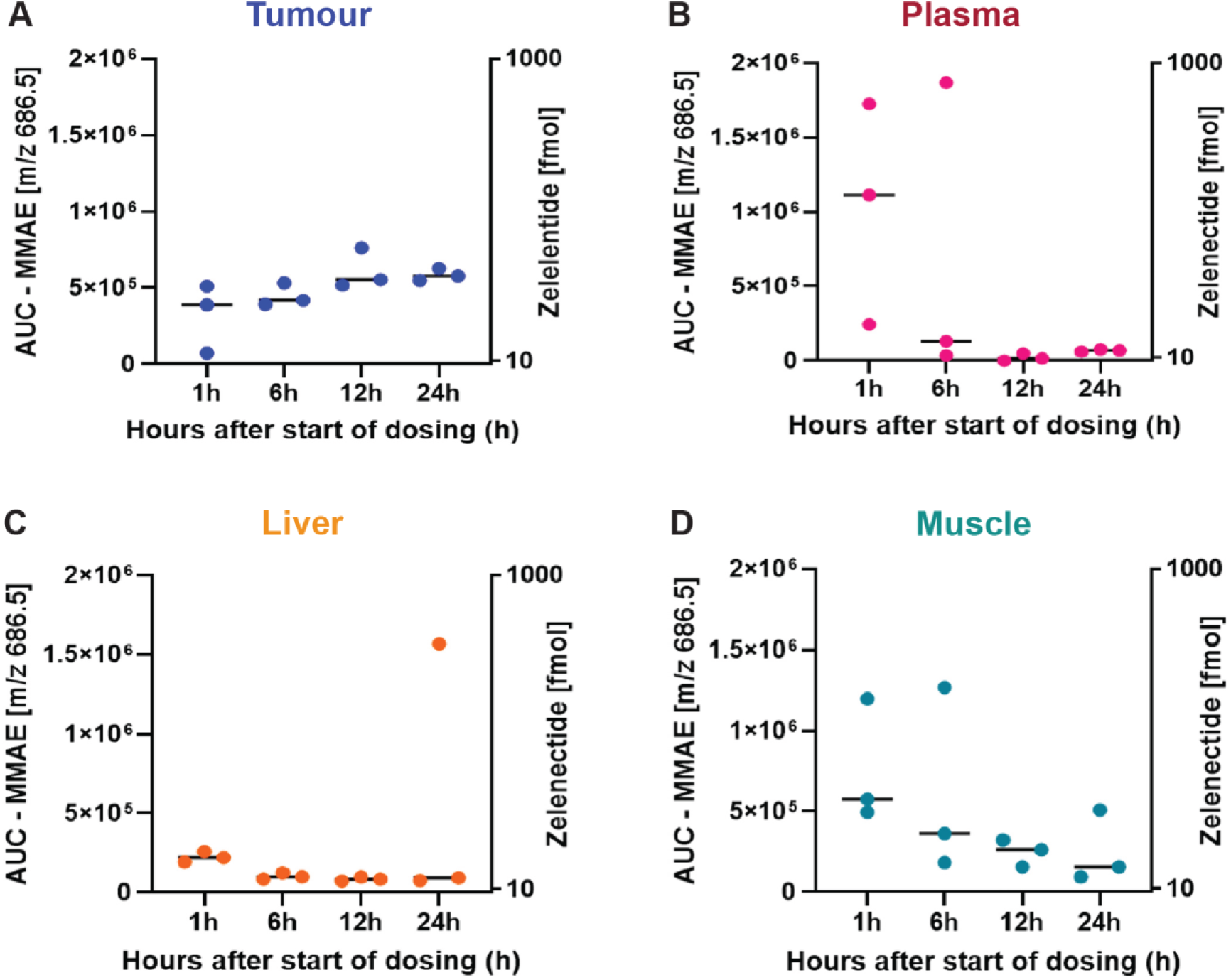
Pharmacokinetic (PK) profile of released MMAE following administration of zelenectide pevedotin across four tissues in the mouse xenograft model reveals higher retention in tumour tissue. **(A)** Tumour tissue exhibited progressively increasing MMAE levels between 1 and 12 h, followed by sustained signal intensity at 24 h, suggesting slow accumulation in the tumour. **(B)** Plasma MMAE levels were highest during the early post-dose period and showed an overall decline over time despite substantial inter-animal variability, consistent with systemic clearance. **(C)** Liver tissue showed an overall reduction in MMAE abundance throughout the study, with one biological replicate at 24 h displaying substantially higher levels than the remaining samples, which may reflect inter-animal variability in hepatic uptake, metabolism or biliary excretion. **(D)** Muscle tissue exhibited transient MMAE exposure without evidence of sustained accumulation, indicating limited tissue retention. The x-axis represents post-dose sampling time (1, 6, 12, and 24 h). The left y-axis shows LC-PRM signal intensity expressed as peak area (AUC) of the quantifier transition (*m/z* 686.5) measured in the *in vivo* samples. The right y-axis provides a calibration reference scale (1-1000 fmol per 1 μL injection) derived from the external calibration curve generated using intact zelenectide pevedotin standards and is shown for reference only. Each data point represents an individual biological replicate (n = 3 per timepoint).

Measured abundances of MMAE in tumour, plasma, liver, and muscle tissues were evaluated against the *in vitro*-determined LOD (4.76 fmol on-column) and LOQ (14.41 fmol on-column) for the quantifier ion. Values above the LOQ were considered fully quantifiable, whereas signals between LOD and LOQ were defined as detectable but below the reliable quantitation limit, and signals below the LOD were considered undetectable or non-reproducible. For example, tumour tissue amounts 1h post-administration ranged from 9.1 to 183.9 fmol on-column, with the majority falling above the LOQ. Similarly, plasma and liver samples at later time points frequently declined below the LOQ, with several measurements approaching or dropping below the LOD, reflecting rapid systemic clearance and tissue-specific clearance.

Quantitative pharmacokinetic profiles were generated using the external calibration curve constructed from pure zelenectide pevedotin standards (10-1000 nM; 10-1000 fmol on-column). Peak areas obtained from the dominant PRM transition (*m/z* 686.5) were interpreted using an external calibration curve generated from intact zelenectide pevedotin standards, allowing semi-quantitative estimation of MMAE-associated signal across tissues and timepoints. A secondary calibration scale (1–1000 fmol per 1 μL injection, with marked LOD = 4.76 fmol and LOQ = 14.41 fmol thresholds) is provided in Figure 3 to facilitate biological interpretation of the measured LC-PRM signal intensities.

The pharmacokinetic profiles of MMAE in tumour, plasma, liver, and muscle tissues were assessed at 1, 6, 12, and 24 h following administration of zelenectide pevedotin (**Figures 3A-D**). For each timepoint, tissue and plasma samples were collected from the same individual animals (n = 3), enabling paired comparisons between biological compartments. Among the analysed tissues, tumour samples exhibited the most sustained MMAE levels over time.

Tumour MMAE concentrations increased progressively throughout the study period (**Figure 3A**). At 1h post-dose, tumour MMAE concentrations ranged from corresponding to 9.1 to 183.9 fmol on-column (mean: 109.5 fmol), reflecting substantial inter-animal biological variability. Tumour MMAE concentrations increased further at 6 h post-dose (136.9-192.7 fmol; mean: 159.1 fmol) and reached the highest mean level at 12 h post-dose (mean: 224.2 fmol; range: 184.5-284.5 fmol). Thereafter, tumour MMAE levels remained relatively stable between 12 and 24 h, suggesting a plateau phase. At 24 h post-dose, tumour MMAE concentrations remained high (199.2-230.8 fmol; mean: 213.7 fmol), indicating sustained tumour-associated MMAE signal throughout the observation period. This accumulation profile is consistent with sustained tumour-associated MMAE signal, while the plateau observed between 12 and 24 h suggests sustained tumour-associated payload exposure over the observation period.

In contrast, plasma concentrations of MMAE demonstrated a generally declining profile following administration (**Figure 3B**). Plasma levels were obtained from the same mice as tissue samples, allowing for paired comparisons. However, due to the limited sample size and high inter-sample variability, no consistent relationship between plasma concentrations and tissue or tumour uptake could be established. Observed differences between individual animals likely reflect biological variability in drug distribution and clearance.

At 1h post-dose, plasma MMAE levels varied markedly, ranging from approximately 77.3 to 665.5 fmol (mean: 380.9 fmol), reflecting initial systemic exposure and substantial inter-animal variability. By 6 h, mean plasma MMAE levels had declined to 240.7 fmol, although substantial inter-animal variability remained, with individual values ranging from below the LOD to 695.3 fmol. At 12 and 24 h, MMAE levels were below the LOD in all samples, consistent with declining circulating MMAE levels over time.

Liver tissue exhibited an overall decline in MMAE abundance over time (**Figure 3C**). At 1h post-dose, concentrations ranged from 57.0 to 83.6 fmol (mean: 69.7 fmol), indicating substantial initial hepatic exposure. Mean levels decreased at 6 and 12 h (21.0 and 14.2 fmol, respectively). At 24 h, two biological replicates remained at low abundance (10.4 and 16.9 fmol), whereas one sample displayed substantially higher abundance (575.4 fmol). This single high-abundance value resulted in a mean of 200.9 fmol at 24 h and represents a marked inter-animal deviation from the other two replicates. Overall, liver samples showed no evidence of consistent MMAE accumulation over the study period.

Muscle tissue exhibited relatively low MMAE abundance throughout the study period, with no evidence of progressive accumulation (**Figure 3D**). At 1 h post-dose, MMAE levels ranged from 176.1 to 453.0 fmol (mean: 278.9 fmol), followed by a decline at 6 and 12 h (mean: 216.0 and 79.1 fmol, respectively). Although concentrations at 24 h varied between biological replicates (range: 17.8-181.6 fmol), no consistent increase or prolonged retention was observed. Overall, these findings indicate transient MMAE exposure in muscle tissue compared with the sustained signal observed in tumour tissue. Overall, tumour tissue exhibited a progressive increase in MMAE abundance between 1 and 12 h, followed by sustained retention at 24 h. In contrast, plasma, liver, and muscle tissues generally showed declining MMAE levels over time, although substantial inter-animal variability remained evident in individual samples. These findings demonstrate distinct tissue-specific MMAE pharmacokinetic profiles following administration of zelenectide pevedotin, with the most sustained signal observed in tumour tissue.

Collectively, these data demonstrate distinct tissue-specific pharmacokinetic profiles of released MMAE following administration of zelenectide pevedotin. Tumour tissue exhibited progressive accumulation and sustained retention throughout the 24 h observation period, whereas plasma, liver and muscle generally showed declining analyte levels over time despite inter-animal variability. Relative tissue-specific distribution patterns are further illustrated by hierarchical clustering of z-score-normalised LC-PRM signal intensities (**Figure 4**). Z-score normalisation was applied exclusively for visualisation and was not used for quantitative comparisons or statistical analyses.

**Figure 4.**
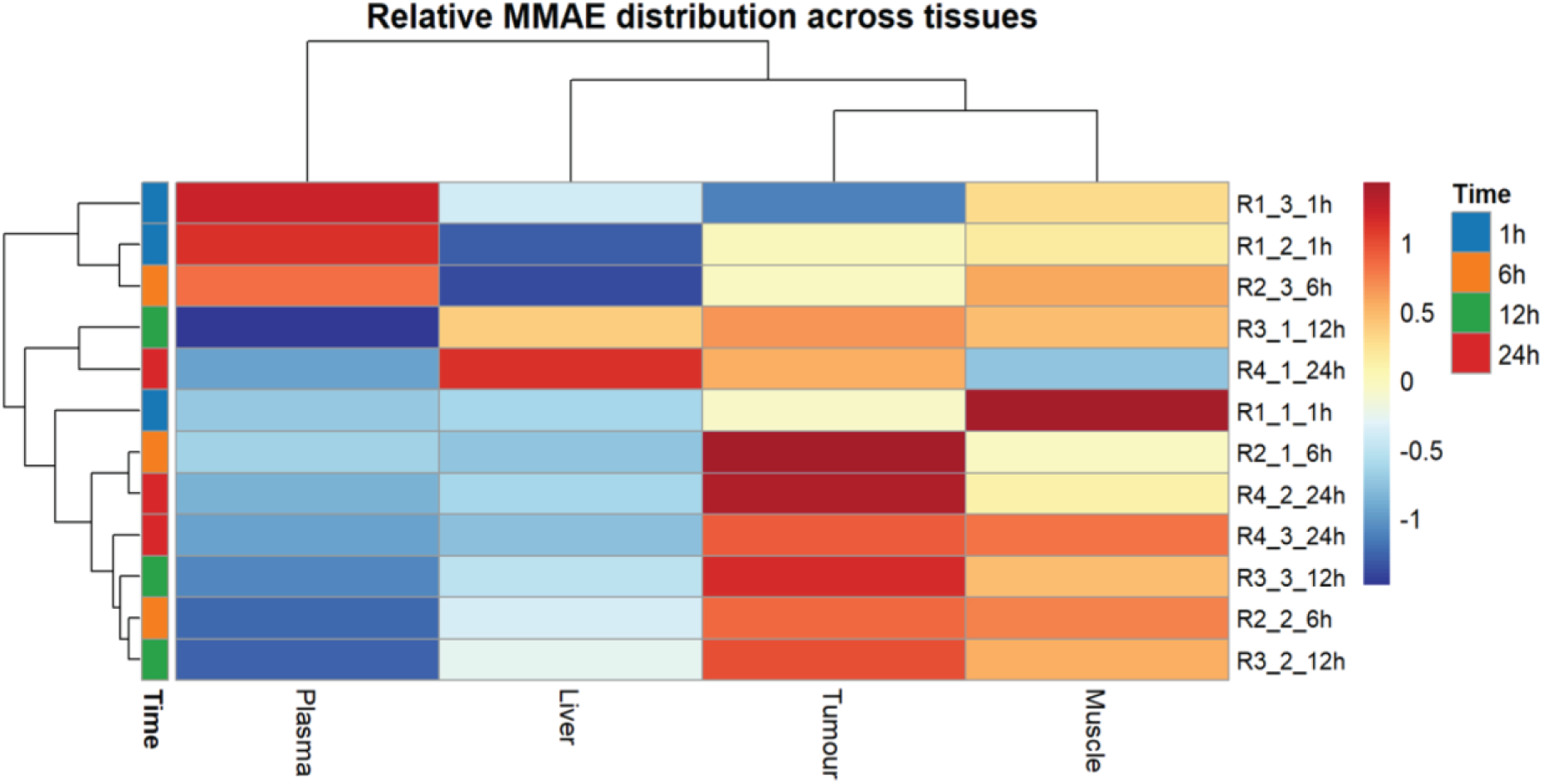
Heatmap of MMAE signal intensities (row-wise z-score normalised per sample) across plasma, tumour, liver, and muscle samples collected at 1, 6, 12, and 24 h post-administration of zelenectide pevedotin. Each row represents an individual biological replicate from xenograft-bearing mice, and columns correspond to tissue types. Relative MMAE abundance was estimated using raw area under the curve (AUC) values of the dominant fragment ion (*m/z* 686.5) acquired via LC-PRM. Hierarchical clustering was performed using Euclidean distance and complete linkage to visualise similarities in tissue-distribution profiles among individual samples. The heatmap highlights relative differences between tissue types within each sample, with tumour samples showing relatively sustained MMAE-associated signal at later timepoints compared with the other tissue compartments. These results support the tissue-specific pharmacokinetics of zelenectide pevedotin and its selective retention within the tumour microenvironment. Z-score normalisation is used here solely to visualise relative tissue-specific distribution patterns and was not used for quantitative comparisons.

### Intra-tumour spatial distribution of zelenectide pevedotin (MMAE) assessed by LCM-LC-PRM

To investigate the intra-tumoural distribution of released MMAE following administration of zelenectide pevedotin, laser capture microdissection (LCM) was combined with LC-PRM to quantify released MMAE in tumour cell-rich (TC) and stroma-rich (TS) compartments across four post-administration timepoints (1, 6, 12, and 24 h). Spatial regions were defined based on histomorphological features observed in adjacent H&E-stained sections, enabling compartment-specific microdissection of tumour cell-dense and extracellular matrix-rich areas for downstream quantitative analysis.

To evaluate the potential impact of histological processing on MMAE detectability, cryosections collected at the 12 h post-dose timepoint (identified as the peak of tumour-associated MMAE abundance by bulk LC-PRM analysis) were analysed using adjacent serial sections that were either stained with hematoxylin and eosin (H&E) or left unstained. Both conditions were processed under identical LC-PRM acquisition settings. H&E-stained sections exhibited a marked reduction in MMAE signal intensity for the quantifier ion (*m/z* 686.5) compared with unstained sections, consistent with partial analyte loss or reduced extraction efficiency following histological staining. In addition, reduced signal intensities were observed for several lower-abundance fragment ions, indicating that H&E processing compromises analyte recovery. Consequently, all subsequent LCM-LC-PRM analyses were performed using unstained tissue sections to preserve analyte integrity and maximise analytical sensitivity (**Figure S6**).

Laser capture microdissection (LCM) was used to isolate defined intra-tumoural regions of interest (ROIs) based on morphological features identified in adjacent H&E-stained sections. Tumour cell-rich (TC) regions were characterised by high cellular density and compact nuclear architecture, whereas stroma-rich (TS) regions were defined by increased extracellular matrix content and reduced tumour cell density. This approach enabled spatially resolved sampling of distinct tumour compartments for downstream LC-PRM analysis of MMAE distribution (**Figure 5A-C**).

**Figure 5.**
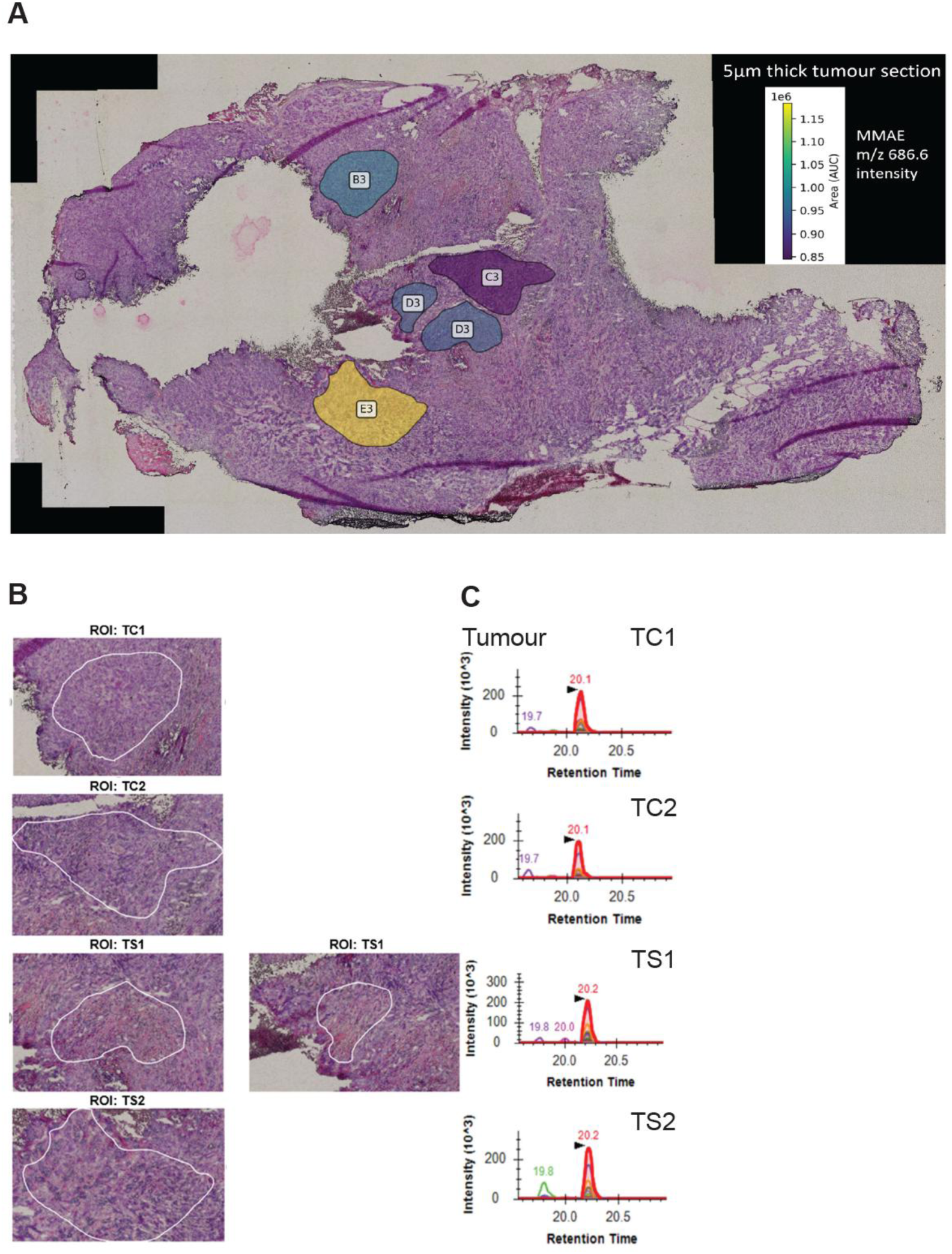
Spatial distribution of zelenectide pevedotin (MMAE) in mouse tumour tissue assessed by LCM-LC-PRM. **(A)** Representative 5 μm-thick H&E-stained tumour section shown for visualisation of tissue morphology. Although all LCM-LC-MS/MS analyses were performed on unstained sections to preserve MMAE integrity, the stained section highlights the regions of interest (ROIs) based on morphological features observed in microscope. **(B)** Magnified view of the four laser-captured tumour regions (TC1, TC2, TS1, TS2), showing spatially distinct tumour cell- and stroma-rich areas selected for LCM. TS1 represents the combination of two adjacent stroma-rich regions for consistency in section size. These structurally distinct subregions were selected based on their morphological differences to enable spatial profiling of drug distribution within the tumour microenvironment. **(C)** Extracted chromatograms from the LCM-PRM analysis, processed using Skyline, following the same analytical approach as the initial LC-PRM analysis of whole tissue. The chromatograms demonstrate reproducible detection of MMAE-associated fragment ions across the microdissected regions. ROI labels denote tissue category (TC = tumour cell-rich; TS = stroma-rich).

The LC-PRM method enabled consistent detection of MMAE-related fragment ions in microscale LCM-derived samples, allowing spatially resolved quantification of MMAE-associated signal across tumour compartments. Chromatographic alignment and co-elution of multiple PRM transitions confirmed analytical specificity and robustness of detection (**Figures S7-S8**). Spatially resolved LC-PRM analysis revealed heterogeneous MMAE-associated signal distribution across TC and TS regions over time, broadly consistent with the temporal patterns observed in bulk tissue analyses (**Supplementary Table A1**).

Measured abundances of MMAE in TC and TS regions were compared to the assay-specific limits of detection (LOD: 1.50 fmol on-column) and quantitation (LOQ: 4.54 fmol on-column), derived according to ICH Q2(R2) guidelines using the standard deviation of the mean responses across technical triplicate injections (*n*=3). Signal levels above 4.54 fmol on-column were considered fully quantifiable, with precision verified to satisfy % *CV* ≤ 20% acceptance criteria (including low-input 1–3 fmol standards displaying % *CV* ≤ 16%). Measured values between 1.50 and 4.54 fmol on-column were defined as detectable but below the reliable quantitation limit, whereas signals below 1.50 fmol on-column were considered undetectable.

Measured MMAE signal intensities were compared across TC and TS compartments at four post-administration timepoints (1, 6, 12, and 24 h). At 1 h, both compartments exhibited high and heterogeneous signal intensities, with TC values ranging from 1.9-3.6 × 10⁶ AUC and TS values ranging from 1.4-3.5 × 10⁶ AUC. At 6 h, increased variability was observed in both compartments, with several TS regions displaying signal intensities comparable to or exceeding those measured in TC regions. By 12 h, overall MMAE signal intensity had decreased, although substantial biological variability persisted between individual samples. At 24 h post-dose, MMAE remained readily detectable in both compartments, with overlapping signal distributions between TC and TS regions.

Overall, these data demonstrate a heterogeneous spatial distribution of MMAE within tumour tissue over time, with both tumour cell-rich and stromal regions contributing to MMAE retention rather than a strictly compartment-exclusive accumulation pattern.

Estimated analyte amounts were derived using the experimentally generated external calibration curve constructed from intact zelenectide pevedotin standards (1-1000 nM, corresponding to 1-1000 fmol on-column for a 1 μL injection; *n*=3 technical triplicates per point). Applying the mean-response linear regression equation (*y*=40534x + 180244; *R^2^*= 0.9992) converted PRM peak areas into estimated MMAE-associated amounts on-column. At 1 h post-dose, TC regions ranged from 42.5 to 85.3 (mean: 70.6), whereas TS regions ranged from 31.5 to 81.4 (mean: 59.8). At 6 h, TC regions ranged from 5.3 to 104.0 (mean: 66.0), while TS regions ranged from 35.2 to 113.0 (mean: 83.7), with higher mean values observed in TS regions than TC regions at this timepoint. By 12 h, values had declined in both compartments. TC regions ranged from 5.1 to 86.1 (mean: 50.5), whereas TS regions ranged from 4.8 to 58.4 (mean: 23.1; with all values remaining strictly above the LOQ threshold of 4.54 fmol). At 24 h post-dose, MMAE remained detectable in all analysed regions, with TC values ranging from 14.3 to 56.8 (mean: 37.9) and TS values ranging from 4.8 to 58.7 (mean: 36.7) fmol on-column.

Overall, tumour cell-rich regions exhibited higher mean MMAE levels at 1, 12, and 24 h, whereas stromal regions displayed higher mean levels at the 6 h. Although substantial variability was observed between individual micro-dissected regions, these data suggest temporal differences in MMAE distribution between tumour cell-rich and stromal compartments. The transient increase in stromal MMAE levels at 6 h distinguished this timepoint from the overall temporal trend observed in TC regions. These findings should, however, be interpreted cautiously given the biological variability observed between individual tumours.

The temporal patterns of MMAE-associated signal across TC and TS regions were further visualised using Z-score-normalised LCM-LC-PRM intensities (**Figure 6A**), with samples displayed according to post-dose timepoint (1, 6, 12, and 24 h). The heatmap highlights the heterogeneous temporal distribution of MMAE-associated signal across the two tumour compartments, including the relatively higher stromal signal observed at 6 h followed by a marked reduction at 12 h. In contrast, TC regions showed comparatively higher relative signal at the earlier and later timepoints. A representative H&E-stained tumour section illustrating compartment selection prior to microdissection is shown in **Figure 6B**. Calibration curves generated from pure zelenectide pevedotin standards (1-1000 fmol on-column) were used to convert PRM peak areas into estimated MMAE amounts for downstream spatial pharmacokinetic analysis (**Figure 6C**).

**Figure 6.**
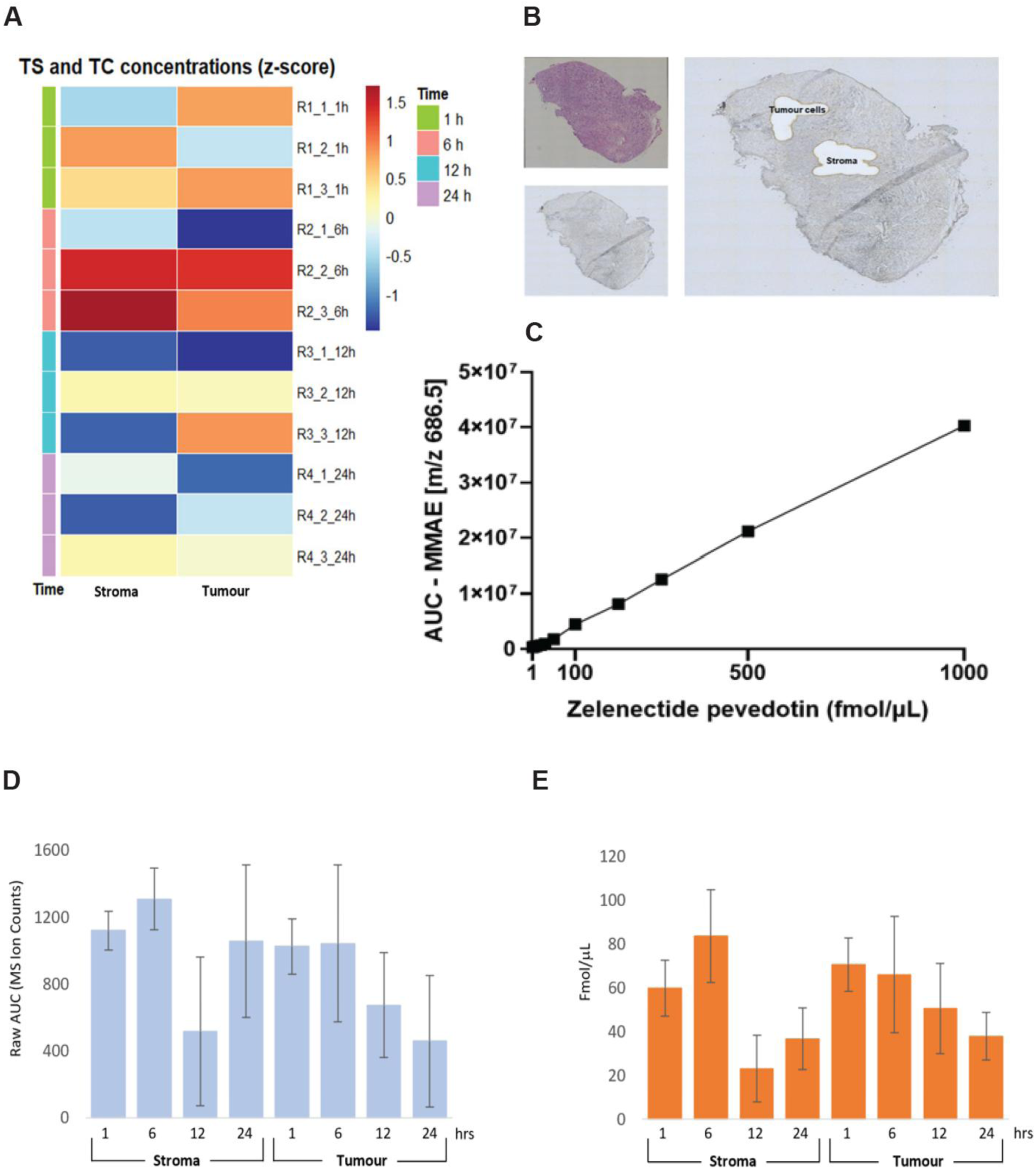
Temporal zelenectide pevedotin (MMAE) and NECTIN-4 abundance profiles in mouse tumour and stroma assessed by LCM-LC-PRM. (A) Heatmap showing Z-score-normalised MMAE abundances measured from the characteristic fragment ion (*m/z* 686.5) in tumour-cell-rich (TC) and tumour-stroma-rich (TS) micro-dissected regions across all post-dose timepoints. Samples are displayed according to post-dose timepoint (1, 6, 12, and 24 h), allowing visualisation of temporal changes in relative MMAE-associated signal. Z-scores were calculated independently for each tissue compartment from raw area under the curve (AUC) values. Each row represents a micro-dissected region from an individual mouse (n = 3 biological replicates per timepoint; four timepoints analysed). Warmer colours indicate higher relative signal intensities. The heatmap highlights the heterogeneous temporal distribution of MMAE-associated signal across TC and TS compartments, including relatively higher stromal signal at 6 h followed by reduced relative signal at later timepoints. **(Β)** Representative 5-µm-thick xenograft tumour section (10× magnification) collected 1 h post-dose, illustrating the regions selected for laser microdissection. The H&E-stained section (top left) was used solely for morphological visualisation and identification of TC and TS regions. The corresponding unstained section (bottom left) was used for the actual laser microdissection. The right panel shows the LCM-captured areas with annotated TC and TS compartments, which served as the basis for compartment-resolved quantitation. TC and TS boundaries were manually defined based on histomorphological criteria and guided the compartment-specific sampling used for downstream PRM-based MMAE quantification. **(C)** Calibration curve of pure zelenectide pevedotin standards. A linear calibration curve was generated by plotting mean AUC (n=3 technical triplicates) against known zelenectide pevedotin amounts (1-1000 fmol on-column; *R^2^* = 0.9992, LOD = 1.50 fmol, LOQ = 4.54 fmol on-column). The resulting regression equation (y = 40534x + 180244) was used to convert PRM peak areas measured in micro-dissected TC and TS regions into estimated analyte amounts (fmol on-column). (**D - E**) Comparison of time-resolved NECTIN-4 protein (light blue) and estimated MMAE (orange) abundance profiles in stroma and tumour tissue sections. 1, 6, 12 and 24 h time points are shown for tissue areas as shown in (**B**). Error bars represent triplicate measurements by LC-MS are presented as mean ± SD from three biological replicates (n = 3). Matched spatial analysis demonstrated a moderate positive trend towards spatial co-localization between NECTIN-4 and MMAE abundance (Pearson r = 0.42, *p* = 0.09; Spearman *ρ* = 0.33).

To further investigate whether spatial NECTIN-4 abundance was associated with MMAE distribution, DIA-based spatial proteomic analysis was performed on matched LCM-isolated tumour cell-rich (TC) and stroma-rich (TS) regions. DIA-derived NECTIN-4 abundances were subsequently compared with LCM-LC-PRM measurements of MMAE obtained from corresponding regions and timepoints (**Figure 6D-E**). Both NECTIN-4 and MMAE abundances displayed parallel temporal trends, with higher relative signals at early timepoints followed by a sharp concurrent reduction in stromal regions between 6 and 12 h post-injection. Bivariate spatial correlation analysis restricted to micro-regions with quantifiable protein signal (*n* = 17 out of 24 analysed regions (70.8%); non-detectable (ND) values were excluded to avoid imputation bias) revealed a moderate positive trend towards spatial co-localization between local NECTIN-4 abundance and MMAE levels (Pearson *r* = 0.42, *p* = 0.09; Spearman *ρ* = 0.33). Although this trend did not reach formal statistical significance at the conventional *α* = 0.05 threshold, the observed directional association is compatible with a potential contribution of NECTIN-4 to local MMAE distribution, although the present data do not establish a causal relationship or directly resolve cellular uptake or payload-release mechanisms. Co-localisation of detectable NECTIN-4 signal with higher MMAE levels was particularly prominent in tumour cell-rich regions at early timepoints, consistent with an association between local target abundance and MMAE distribution.

Collectively, these findings suggest that local NECTIN-4 abundance may contribute to intra-tumoural MMAE localization following zelenectide pevedotin administration, while additional microenvironmental factors are likely to influence overall payload distribution across distinct tissue micro-compartments.

## Discussion

Here, we explored LCM-LC-PRM for spatial pharmacokinetic analyses of a bicyclic drug conjugate (BDC), zelenectide pevedotin, at microscale resolution *in vivo*. The observed intra-tumoural heterogeneity highlights the importance of spatial profiling in accurately characterising the biodistribution of BDCs.

Targeted delivery of cytotoxic agents is a critical parameter of precision oncology. However, its translation to solid tumours remains challenging due to intratumoural heterogeneity and physical barriers. By optimising a combination of laser capture microdissection (LCM) and highly sensitive parallel reaction monitoring (PRM) mass spectrometry, we achieved spatially resolved detection and estimated quantification of MMAE distribution within histologically defined tumour compartments and bulk non-tumour tissues (**Figure S9**).

To support this spatial characterisation, we performed two complementary LC-PRM experiments. Bulk LC-PRM analysis revealed detectable MMAE in all analysed tissues at 1 h post-dose. Tumour tissue exhibited a progressive increase in MMAE abundance between 1 and 12 h, followed by sustained retention at 24 h, whereas plasma, liver and muscle generally showed declining signals over time (**Figure 3**). These distinct temporal profiles suggest prolonged retention of MMAE within tumour tissue relative to systemic circulation and non-tumour tissues. Although substantial inter-individual variability was observed, particularly in plasma and liver, the overall tissue distribution pattern was consistent with sustained tumour-associated MMAE abundance over the 24 h observation period.

Spatially resolved analysis of LCM-isolated tissue compartments showed distinct kinetics. At 1 h, tumour cell-rich (TC) regions showed a higher mean MMAE abundance than stroma-rich (TS) regions, whereas the relationship was reversed at 6 h, when TS regions exhibited the highest mean signal. By 12 h, MMAE abundance was again higher in TC regions, while at 24 h comparable levels remained detectable in both compartments (Figure 6). This divergence underscores the dynamic redistribution of the toxin cargo across tumour microenvironments, which cannot be resolved by bulk measurements. The transiently higher MMAE abundance in stromal regions at 6 h may reflect redistribution of released toxin from initially exposed tumour regions into surrounding extracellular compartments, although the present data cannot directly distinguish diffusion from other processes contributing to this spatial pattern.

Our results demonstrate that an MMAE-derived signal associated with zelenectide pevedotin exhibits selective and sustained accumulation within tumour tissues, whereas plasma, liver and muscle showed transient exposure followed by declining analyte abundance over time. This is consistent with the molecular design principles of BDCs, which enable rapid tumour penetration, efficient enzymatic cleavage, and toxin retention in the tumour microenvironment (TME)[12,26,27]. Interestingly, we did not detect intact BDC nor the bicyclic peptide moiety in the tissue context, suggesting rapid clearance of these molecular components. In contrast, the prolonged tumour-associated MMAE signal observed after the decline of circulating MMAE is consistent with retention of the MMAE-associated payload signal within the tumour microenvironment. However, because intact zelenectide pevedotin was not reliably quantifiable in the in vivo samples, the present study cannot directly define the relative contributions of intact BDC delivery, local cleavage, and subsequent MMAE redistribution. These findings support the hypothesis that BDCs, unlike ADCs, may undergo distinct processing pathways that do not depend exclusively on intracellular trafficking and lysosomal degradation, thereby potentially bypassing common resistance mechanisms associated with altered internalisation or endosomal escape[28,29].

Spatial resolution achieved via LCM enabled us to dissect the tumour into distinct regions-tumour cell-rich (TC) versus tumour stroma-rich (TS) and quantify drug distribution across compartments. Rather than demonstrating a fixed preferential accumulation in one compartment, our results revealed a dynamic and heterogeneous spatial distribution of MMAE. TC regions showed higher mean abundance at 1 h, TS regions showed higher mean abundance at 6 h, and both compartments retained readily detectable MMAE at 24 h. A transient increase in stromal MMAE abundance at 6 h may reflect redistribution through the tumour microenvironment, potentially influenced by vascular accessibility, extracellular matrix composition, and local diffusion dynamics. These spatial gradients are critical for understanding both efficacy and toxicity, particularly in the context of heterogeneous antigen expression and stromal shielding, factors that frequently limit the performance of antibody-based modalities such as ADCs, but are handled differently by peptide-based systems such as BDCs[30–32].

Molecular targets, such as NECTIN-4, could, in principle, influence regional drug distribution in certain tumour types. The observed MMAE distribution profile is broadly consistent with reports that NECTIN-4 expression increases with tumour progression across several cancer types [33–35]. While untargeted proteomic analysis of bulk xenografts using DDA did not detect NECTIN-4 - in line with public databases (Human Protein Atlas) showing moderate-to-low expression in lung cancer lineages such as NCI-H322 - high-sensitivity DIA-based spatial proteomics enabled robust quantification across micro-dissected subregions. To investigate whether intra-tumoural MMAE localization was associated with local target abundance, we integrated spatial DIA-derived NECTIN-4 measurements with matched LCM-LC-PRM MMAE quantification across TC and TS regions (**Figure 6**). Bivariate analysis across quantifiable micro-regions revealed a moderate positive trend towards spatial co-localization between local NECTIN-4 abundance and MMAE levels, indicating an association between NECTIN-4 abundance and local MMAE distribution. Co-localisation of detectable NECTIN-4 signal with higher MMAE levels was observed in tumour cell-rich compartments at early timepoints, consistent with an association between local target abundance and MMAE distribution. Importantly, the present study does not directly assess zelenectide pevedotin binding to NECTIN-4, cellular internalisation, intracellular trafficking, or the mechanism of MMAE release; these processes therefore remain mechanistic hypotheses requiring dedicated experimental investigation. Subsequently, MMAE was preferentially retained in tumour cell relative to stroma fractions at later time points (**Figure 6D-E)**. Beyond target expression alone, this pharmacokinetic profile may be modulated by local physiological factors, including tissue architecture, vascular accessibility, extracellular matrix composition, and diffusion dynamics within the tumour microenvironment[36].

Beyond tumour tissue, our data revealed tissue-specific differences in MMAE abundance, consistent with previous reports[28,37,38]. We observed some inter-individual variability, particularly in liver samples. Liver MMAE abundance decreased between 1 and 12 h, although substantial variability was observed at 24 h, driven by one biological replicate with markedly elevated signal. Plasma samples exhibited the expected decline in MMAE abundance over time, consistent with systemic clearance, whereas muscle tissue showed comparatively low signal intensities and no evidence of sustained accumulation throughout the study, indicating limited persistence in this non-target tissue. Collectively, these findings support preferential tumour retention of MMAE while demonstrating that LC-PRM enables sensitive and reproducible assessment of tissue-specific MMAE distribution profiles multiple biological matrices. Interpretation of the hepatic findings should, however, remain cautious given the observed biological variability and the complexity of MMAE metabolism, which extends beyond the scope of the present study.

Further pharmacokinetic profiling beyond 24 h would be valuable to determine whether MMAE levels remain elevated or eventually decline in tumour tissue, which may inform dosing strategies.

While the whole-tumour PRM analysis provided valuable temporal insights into zelenectide pevedotin pharmacokinetics, demonstrating a clear plateau in MMAE accumulation between 12 and 24 h post-administration, this bulk approach inherently lacks spatial resolution and may obscure key regional differences in distribution. In contrast, the LCM-based approach uncovered dynamic region-specific distribution patterns, including the transiently higher MMAE abundance observed in stromal regions at 6 h and the comparable retention observed in both compartments at 24 h. This comparison underscores the limitations of conventional bulk methods and highlights the value of spatially resolved mass spectrometry in preclinical pharmacology. The present findings further underscore the importance of detailed tissue-level pharmacokinetic profiling, particularly for payloads such as MMAE that are shared across different drug platforms[28,37,38].

The methodological approach we present, combining LCM with targeted LC-PRM, provides a versatile and scalable workflow for spatial drug quantification. Unlike traditional LC-MS/MS, our platform enables both molecular specificity and histological compartmentalisation. This makes it especially well-suited to next-generation therapeutics such as BDCs (a structurally constrained subclass of peptide-drug conjugates, PDCs), and proteolysis-targeting chimeras (PROTACs), which may act extracellularly or engage with complex tissue microenvironments[26,39–41].

Nevertheless, some limitations warrant consideration. The quantification of payload distribution was limited to static histological snapshots at defined timepoints and the integration of dynamic imaging methods, such as MALDI imaging or fluorescence-based probes, could complement our findings. Additionally, while murine xenograft models offer controlled conditions and reproducibility for spatial analysis, they may not fully recapitulate the complexity of human tumour vasculature, antigen heterogeneity, or immune infiltration. The relatively small number of biological replicates at each timepoint (n = 3) also limits the statistical power to resolve differences between tumour compartments and to distinguish biological variability from treatment-related spatial effects.

Furthermore, in the absence of an isotopically labeled internal standard for free MMAE, payload abundance was estimated using an external calibration curve generated from pure reference standards; therefore, the reported values represent estimated zelenectide pevedotin-equivalent amounts of payload on-column rather than absolute free MMAE concentrations. Validation in clinical samples will therefore be essential.

Taken together, this study establishes a robust spatial pharmacology workflow that enables precise, high-resolution assessment of zelenectide pevedotin, a peptide-drug conjugate-released toxin across histologically distinct tumour compartments. The results reveal critical spatial pharmacokinetic heterogeneity not captured by bulk approaches and highlight the importance of compartment-specific profiling in understanding drug behaviour in solid tumours. Importantly, the dynamic differences observed between tumour cell-rich and stromal regions demonstrate that tumour exposure is not spatially uniform and changes over time following drug administration. The observed sustained tumour-associated payload signal highlights the potential of BDCs as a strategy for achieving prolonged tumour exposure, while motivating further investigation of the cellular and molecular mechanisms underlying payload distribution and retention.

## Materials and Methods

### Chemicals and reagents

Analytical standard zelenectide pevedotin (MW: 4173.85 g/mol), provided by Bicycle Therapeutics, was used to prepare a primary stock solution. Reagents including RIPA Lysis and Extraction Buffer, sodium dodecyl sulphate (SDS) 20% solution (RNase-free), Pierce™ BCA Protein Assay Kit, dithiothreitol (Pierce™ DTT), urea, Tris(hydroxymethyl)aminomethane hydrochloride (Tris-HCl), and n-Dodecyl-β-D-maltoside (DDM) were obtained from Thermo Fisher Scientific. HPLC-grade acetonitrile (ACN), formic acid (FA), trifluoroacetic acid (TFA), dimethyl sulphoxide (DMSO), ultrapure water, chloroform (CHCl₃), and methanol (MeOH) were purchased from Thermo Fisher Scientific. Iodoacetamide (IAA) and triethylammonium bicarbonate buffer (TEAB) were obtained from Sigma-Aldrich, while sequencing-grade modified trypsin and Trypsin/LysC Mix were sourced from Promega. Protease and phosphatase inhibitors, including cOmplete™ Mini EDTA-free Protease Inhibitor Cocktail and PhosSTOP (both from Roche), as well as Benzonase® Nuclease (Millipore), were also used. For histological processing, denatured alcohol, Gill’s III haematoxylin, eosin, and hydrochloric acid were purchased from Sigma-Aldrich.

### Standard compound preparation

A primary stock solution of zelenectide pevedotin was prepared by dissolving 10.32 mg of zelenectide pevedotin in 1 mL of DMSO, yielding a 1% (w/v) solution (10.32 mg/mL, corresponding to approximately 2.4 mM). The solution was vortexed for 2 minutes and sonicated for 5 minutes to ensure complete dissolution of the analyte. The resulting stock solution was aliquoted into 100 µL single-use portions to avoid repeated freeze-thaw cycles and stored at -80 °C.

A single aliquot was subsequently thawed and diluted with DMSO to prepare a working solution at a final concentration of 1 mM. Additional solutions for mass spectrometry analysis were prepared by serial dilution of the working solution in a mixture of 5% DMSO and 5% FA to obtain the desired final concentrations. Where necessary, the volume of the stock solution used for dilution was adjusted to achieve the desired concentration.

### Mass spectrometry method development

In order to detect zelenectide pevedotin-derived MS/MS signatures in complex biological matrices (e.g., mouse tissue and plasma lysates), a two-phase mass spectrometry method development strategy was implemented. First, data-dependent acquisition (DDA) was initially employed to analyse the pure zelenectide pevedotin reference compound. This step aimed to confirm compound detection, determine charge states, optimise fragmentation conditions, and establish accurate retention time (RT) under high-resolution conditions. Since DDA alone was not sufficiently sensitive for detecting zelenectide pevedotin in complex *in vivo* samples, these optimised parameters were subsequently used to configure a parallel reaction monitoring (PRM) method, which was applied to biological matrices to enhance specificity and sensitivity of detection as well as accurate quantitation.

All mass spectrometry experiments were performed on an UltiMate™ 3000 UHPLC system (Thermo Fisher Scientific) coupled to a nano-electrospray ion source on a Q Exactive™ Hybrid Quadrupole-Orbitrap™ Mass Spectrometer (Thermo Fisher Scientific). Samples were initially loaded onto a trapping column (Acclaim™ PepMap™ 100 C18 HPLC Column; 300 µm × 5 mm, 5 µm particle size, Thermo Fisher Scientific) and subsequently separated on an analytical column (EASY-Spray™ PepMap™ RSLC C18, 2 µm, 100 Å, 75 µm × 50 cm; Thermo Fisher Scientific). Chromatographic gradients and MS acquisition parameters are detailed for each mode below.

### Data dependent mass spectrometry analysis

In the initial method development phase, the zelenectide pevedotin reference standard was analysed using data-dependent analysis (DDA) with a 60-minute gradient to confirm compound detection, determine charge states (+2, +3, +4), and establish accurate RT under high-resolution chromatographic conditions. Normalised collision energy (NCE) was optimised by testing a range of values to enhance fragmentation efficiency. The experimentally determined retention time (RT), optimal normalised collision energy (NCE), and tandem MS/MS spectra were then used to configure the targeted PRM method for screening and identification of the compound, which employed a shorter 35-minute gradient.

Samples were resuspended in 5% FA and 5% DMSO. Peptides were loaded using loading solvent A (0.1% FA in water) at 60 bar, and eluted with a linear gradient over 60 minutes (2% to 60% solvent B [0.1% FA in ACN and 5% DMSO]) at 250 nL/min. Raw data were acquired in Full-MS/ddMS2 mode with the following settings: full MS scan range 380-2200 *m/z*, resolution 70,000, AGC target 3e6, max injection time 100 ms. The 15 most intense ions were selected for HCD fragmentation at 44% NCE, acquiring MS2 spectra at 17,500 resolution, AGC target 1e5, max injection time 128 ms. Charge exclusion was applied to unassigned, 1+ and >8+ charged ions, and dynamic exclusion was set to 7 seconds.

### Untargeted proteomic analysis and MaxQuant processing

Untargeted proteomic analysis was performed on bulk tumour lysates corresponding to the xenograft samples using the optimised MS/MS method that also included PRM-based MMAE quantification. Peptide digests were analysed by data-dependent acquisition (DDA) on a Q Exactive Orbitrap mass spectrometer using a 60-minute linear gradient (1-60% solvent B), as described above.

Raw mass spectrometry (MS) files were processed using MaxQuant (v2.7.5.0) with the integrated Andromeda search engine. Spectra were searched against the UniProt Human and Mus musculus reference proteomes, supplemented with common contaminants. Carbamidomethylation of cysteine was set as a fixed modification, while methionine oxidation and N-terminal acetylation were specified as variable modifications. Trypsin/P was selected as the digestion enzyme, allowing up to two missed cleavages.

Precursor ion mass tolerance was set to 4.5 ppm and fragment ion mass tolerance to 20 ppm. Peptide-spectrum match (PSM), peptide, and protein false discovery rates (FDR) were controlled at 1% using a target-decoy strategy. Protein identification required at least one unique peptide. Label-free quantification (LFQ) was enabled.

### PRM mass spectrometry analysis for MMAE quantitation

Samples were prepared as described above for DDA analysis. Chromatographic separation was performed with a segmented gradient over 35 minutes: solvent B increased from 7% to 60% in 4 minutes, then to 80% in 8 minutes, held for 5 minutes, and returned to 7% for equilibration, at a flow rate of 250 nL/min.

Raw data were acquired in parallel reaction monitoring (PRM) mode on a Q Exactive Orbitrap mass spectrometer at a resolution of 17,500 (MS2), with an AGC target of 1×10⁶, maximum injection time of 100 ms, an isolation window of 1.6 *m/z*, and a fixed first mass of 100 *m/z*. Stepped normalized collision energy (NCE) was applied, with 44% for zelenectide pevedotin and 28% for MMAE. For matrix-containing samples, a lower NCE of 28% was also applied to the remaining proteome to optimize fragmentation and improve sensitivity.

PRM transitions were monitored for the precursor ions listed in **Table S2**, and the most intense fragment ions (daughter ions) for each compound, as indicated in **Figure 1**, were used for quantification. Mouse tissue and plasma lysates were loaded at ∼200 ng per injection, based on quantification using the Pierce BCA Protein Assay Kit (Thermo Fisher Scientific). Quantification was carried out using the best-performing transitions in PRM mode.

### Calibration curve for pure zelenectide pevedotin standards

Two independent calibration strategies were employed corresponding to the two experimental settings used in this study (bulk tissue analysis and LCM-based spatial analysis). The most intense and reproducible fragment ion (*m/z* 686.5) was selected as the quantifier ion, and peak areas (AUCs) were extracted using Skyline and Xcalibur software. These calibration curves were subsequently used to convert PRM peak areas measured in nLC-PRM and LCM-derived samples into estimated analyte amounts (fmol injected on-column). No signal was detected in blank solvent or drug-free tissue matrices, confirming negligible background interference. Due to the high resolution and mass accuracy of the FTMS Orbitrap platform, baseline noise in Extracted Ion Chromatograms was negligible (*S/N → ∞*). Therefore, limits of detection (LOD) and quantitation (LOQ) for both experimental settings were calculated using the ICH Q2(R2) linear regression approach [LOD = 3.3(*σ/S*) and LOQ = 10(*σ/S*)], where *σ* represents the residual standard deviation of the regression and *S* the slope of the calibration curve.

For bulk tissue pharmacokinetic analysis (Experiment 1), an external calibration curve was generated using pure zelenectide pevedotin standards prepared in solvent, spanning nominal concentrations from 1 to 1000 nM (1, 10, 30, 100, 300, and 1000 nM). Standards were analysed by nLC-PRM under the optimised method conditions described above, using a fixed injection volume of 1 µL on column, utilising a single injection per concentration calibration setting for response-concentration conversion in higher-input samples. The calibration curve was generated by plotting the measured peak area of the quantifier ion (*m/z* 686.5) against the nominal zelenectide pevedotin concentration. Linear regression demonstrated excellent linearity across the evaluated concentration range (R² = 0.998), with a slope (*S*) of 2506 and an intercept of 49422. Matrix effects were evaluated using matched concentration series spiked into control liver homogenates and plasma.

For LCM-derived spatial analysis (Experiment 2), a separate external calibration curve was generated using an expanded dilution series of pure zelenectide pevedotin standards (1, 2, 3, 5, 10, 20, 30, 50, 100, 200, 300, 500, and 1000 nM), corresponding to 1-1000 fmol injected on-column with a 1 μL injection volume. Each calibration point was analyzed in technical triplicates to evaluate quantitative precision at low input levels. The peak area of the quantifier ion (*m/z* 686.5) was used for quantification. Mean peak area (AUC), standard deviation (*SD*), and coefficient of variation (*%CV*) were calculated for each calibration point. Linear regression based on the mean AUC values yielded the equation *y* = 40534x + 180244 (*R²* = 0.9992). High analytical precision at low input levels was confirmed, with CV values of 12%, 16%, and 10% for the 1, 2, and 3 nM standards, respectively, validating that the calculated LOQ satisfied the acceptance criterion of (%CV) ≤20%).

The implementation of separate calibration curves reflected necessary differences in sample input, biological matrix complexity, and technical replicates between bulk tissue extracts and LCM-derived microsamples rather than variations in analytical performance.

### Control sample preparation and matrix evaluation

Blank liver, plasma, and muscle samples were obtained from wild-type (WT) female BALB/c mice (n = 3 per tissue type), aged 6-8 weeks and sourced independently from the samples provided by Bicycle Therapeutics. The mice were supplied by the Biomedical Sciences Animal Facilities of the University of Oxford, UK. All procedures complied with the UK Animals (Scientific Procedures) Act.

Tissue homogenisation was performed in two steps. Flash-frozen samples were first pulverised under liquid nitrogen using a porcelain mortar and pestle. The resulting powder was further disrupted to nanoscale particles by bead beating. Approximately 18 mg (±3 mg) of each powdered tissue sample and 20 microliters of plasma were lysed in 300 µL and 180 µL respectively of ice-cold 1× RIPA buffer containing 2% SDS (ThermoFisher Scientific) and a protease inhibitor cocktail (cOmplete™ Mini, EDTA-free, Roche, #11836170001), with zirconia/silica beads (1.0 mm, Thistle Scientific #11079110) using a Precellys® 24 homogeniser. Lysates were centrifuged at 4000 rpm for 10 minutes at 4 °C and supernatants collected.

Protein concentrations were determined using the Pierce™ BCA Protein Assay Kit (Thermo Scientific). Based on protein quantification, aliquots containing 200 µg of total protein from each tissue lysate and 150 µL from each plasma lysate were prepared. These aliquots were used to generate both blank tissue lysate and zelenectide pevedotin-spiked tissue lysate backgrounds. Pure zelenectide pevedotin was spiked into liver and plasma lysates at concentrations of 10, 30, 100, 200, 300, and 1000 fmol/µL, matching the calibration curve to replicate real sample conditions and evaluate potential matrix effects. All samples then underwent identical extraction and processing procedures.

Proteins were precipitated using chloroform/methanol (CHCl₃/MeOH), reduced with 5 mM DTT for 30 minutes at room temperature and alkylated with 20 mM IAA for 30 minutes in the dark. After precipitation, proteins were resuspended in 50 μL of 6 M urea prepared in 100 mM TEAB and diluted to a final urea concentration of <1 M. Digestion was performed with trypsin at a final concentration of 0.2 µg/µL (enzyme-to-protein ratio 1:50, w/w), overnight at 37°C using an Eppendorf ThermoMixer C (400 rpm). Peptides were acidified with 1% TFA, desalted using C18 solid-phase extraction cartridges (Sep-Pak Plus, Waters), and eluted with 80% ACN/0.1% TFA. Samples were dried using a SpeedVac at room temperature and stored at -80 °C until analysis.

Matrix effects were evaluated by comparing the peak area of the quantifier ion (*m/z* 686.5) corresponding to the dominant MMAE-derived fragment of zelenectide pevedotin, between samples spiked prior to extraction and analyte-only solvent standards. This comparison was used to assess ion suppression or ion enhancement arising from endogenous biological matrix components. The concentrations used for matrix effect evaluation matched those used for the bulk calibration curve. Matrix-matched calibration curves were used solely to assess the influence of biological matrices on LC-PRM signal response, whereas the solvent-based external calibration curve was used for quantitative interpretation of the in vivo pharmacokinetic data.

### Animal studies and *in vivo* analysis

Mouse animal work was performed by WuXi AppTec Co., Ltd. (Shanghai, China).

#### Animal model and tumour induction

A total of 23 BALB/c nude female mice, aged 6-8 weeks and weighing approximately 18-22 g, were sourced from Shanghai Lingchang Biotechnology Experimental Animal Co., LTD., and housed in the Animal House Facility of WuXi AppTec Co., Ltd., Shanghai, China. An acclimatised period for seven days was allowed between animal arrival and tumour inoculation. All mice were housed in a specific pathogen-free environment under controlled conditions at 20-26 °C with 40-70% humidity and a 12 h light/dark cycle and in individual ventilation cages (5 mice per cage). Food and water were provided *ad libitum*.

#### Cell-derived xenograft (CDX)

NCI-H322 tumour cells were maintained in a medium supplemented with 10% heat-inactivated fetal bovine serum at 37°C in an atmosphere of 5% CO₂ in air. The cells were routinely subcultured twice weekly, harvested during their exponential growth phase, and counted for tumour inoculation. Ten million (1 × 10^7^) NCI-H322 tumour cells were mixed with 50% matrigel and injected subcutaneously into the right flank of each mouse to establish tumours. The tumour volume was expressed in mm³ using the formula: V = 0.5 × a × b², where *a* and *b* are the long and short diameters of the tumour, respectively. Following tumour inoculation, the animals were monitored daily for signs of morbidity and mortality. Before the commencement of treatment, all animals were weighed, and their tumour volumes were measured.

When tumours reached a volume of approximately 200 mm³, mice were assigned to treatment groups using a randomized block design based on tumour volume to ensure comparable baseline tumour burdens. Zelenectide pevedotin was administered intravenously at a dose of 3 [unit] with a dosing volume of 10 mL/kg body weight.

For the present analysis, samples from 12 treated animals were analysed, comprising three animals at each of four post-dose timepoints (1, 6, 12, and 24 h; *n = 3* biological replicates per timepoint). From each animal, plasma, solid tumour, liver, and skeletal muscle (gastrocnemius) samples were collected for subsequent analysis (see Table S1, Supplementary Methods).

All procedures involving the animal handling, care, and treatment were reviewed and approved by the Institutional Animal Care and Use Committee (IACUC) at WuXi AppTec Co., Ltd. (Shanghai, China), and were conducted in accordance with the regulations of the Association for Assessment and Accreditation of Laboratory Animal Care (AAALAC).

#### Sample collection and preparation

Mice were anaesthetised and sacrificed at 1, 6, 12, and 24 h following treatment administration. From each individual mouse, approximately 200 μl of blood was collected via tail vein sampling into EDTA-coated tubes. Plasma samples were centrifuged at 12,000 x *g* for 5 minutes at 4 °C, and the supernatant was collected and stored at -80 °C until further processing.

Following plasma collection, mice were euthanised and liver, skeletal muscle (gastrocnemius), and solid tumour tissues were harvested from the same animals. All tissues were snap-frozen on dry ice and subsequently shipped to the Target Discovery Institute / Centre for Medicines Discovery, Nuffield Department of Medicine, University of Oxford (Oxford, UK), for further analysis.

Each tissue sample was divided into two portions. One portion was snap-frozen for quantification of zelenectide pevedotin by LC-PRM, while the second portion was embedded in O.C.T. compound. O.C.T.-embedded tumour tissue was used for cryosectioning and subsequent spatial pharmacology analysis using laser capture microdissection (LCM), whereas O.C.T.-embedded liver and muscle tissues were not subjected to LCM analysis. Tumour tissue blocks were acclimatised to −20 °C and mounted onto cryostat chucks using O.C.T. compound. Care was taken to avoid contamination of tissue sections with excess O.C.T. material.

### LC-PRM analysis in multiple tumour tissues

For LC-PRM analysis, proteins and zelenectide pevedotin (MMAE) were extracted from plasma, liver, muscle, and tumour tissues using the same chloroform/methanol precipitation, reduction, alkylation, and digestion workflow described in Section 4. This workflow was originally established for control and matrix evaluation samples, ensuring full procedural consistency across all analytical steps.

Processed lysates were analysed using the optimised PRM acquisition method detailed in Section 3. To minimise carryover and ensure measurement accuracy, a blank/wash injection followed by a high-organic gradient blank was applied between each run. These steps confirmed effective column clearance and eliminated signal contamination.

Quantification of zelenectide pevedotin-derived MMAE in bulk tissue lysates (experiment 1) was performed using an external calibration curve generated from single-injection standards, whereas quantification in LCM-derived spatial subregions (experiment 2) used calibration curves based on triplicate standards and mean AUC values. Quantitation was based on the MMAE-specific PRM transition at *m/z* 686.5, which demonstrated the highest sensitivity and reproducibility.

PRM signal intensities (AUCs) were converted to estimated analyte amounts (fmol on-column) by applying the corresponding external calibration curve equation (y = ax + b), where AUC values were used to calculate the equivalent amount of analyte injected. Because a fixed injection volume of 1 μL was used throughout the study, all reported values are expressed as fmol on-column.

Appendix A Table C reports the raw peak area values (AUCs) of MMAE measured in bulk plasma, tumour, liver, and muscle tissue samples (experiment 1). Appendix A Table D reports the spatial quantification from LCM-derived tumour subregions (experiment 2).

### Spatial Quantification by LCM-LC-PRM

Tumour cryosections (5 µm) were prepared on UV-irradiated Superfrost™ slides (Thermo Fisher Scientific) and PEN (polyethylene naphthalate) membrane slides (Leica Microsystems) and stored at - 80 °C until analysis. For each tumour block, adjacent paired sections were generated, with one section left unstained for laser microdissection and an adjacent section subjected to H&E staining for histological reference.

To assess the impact of histological staining on MMAE detectability, an additional pair of adjacent sections from a representative 12 h post-dose tumour was analysed both stained and unstained. Unstained sections consistently yielded higher MMAE signal intensities, whereas stained sections showed ∼10-fold reduced signal and selective loss of diagnostic fragment ions. Therefore, all LCM-based quantification was performed exclusively on unstained sections, while H&E-stained sections were used only for morphological guidance (**Figure S8**).

H&E staining was performed as follows: sections were fixed in 70% denatured ethanol, hydrated, stained with Gill’s III haematoxylin, differentiated in 0.4% acid alcohol, and counterstained with 0.5% eosin Y solution, with appropriate washing between steps. Stained sections were dehydrated through graded ethanol, air-dried without coverslips, and stored at -80 °C until further analysis.

Laser capture microdissection (LCM) was performed on annotated tumour subregions using a Leica LMD7 with a 10× objective (power 50, aperture 3, speed 17, pulse count 3, final pulse 18, head current 100%, pulse frequency 1046). Two spatially distinct regions-tumour-cell-rich (TC) and stromal (TS)-were isolated per sample, each corresponding to ∼1 mm² tissue area (∼6 nL, ∼900 ng protein). The regions were annotated on images of the stained adjacent sections in QuPath and transferred to the unstained sections on LMD7 with the Qupath-to-LMD[42]software. Microdissected samples were collected into 96-well PCR plates. Acetonitrile was added to the wells and the plates centrifuged at 2000 g for 10 min to collect tissue pieces at the bottom of the wells. Plates vacuum-dried at 40 °C to evaporate acetonitrile and stored at -80 °C.

Protein and zelenectide pevedotin extraction from LCM-derived tissue was performed using a DDM-assisted solubilisation and Trypsin/Lys-C digestion protocol optimised for low-input, O.C.T.-embedded samples. Samples were thawed and equilibrated at room temperature for 30 minutes, followed by brief centrifugation. Protein and zelenectide pevedotin solubilisation and extraction were performed in 20 µL of 0.013% DDM in 50 mM TEAB, with incubation at 95 °C for 90 minutes, followed by cooling to room temperature. Subsequently, sample volumes were normalised to 20 µL with water, and organic denaturation was performed by the addition of 5 µL 60% ACN, followed by incubation at 75°C for 30 minutes and cooling to room temperature. The ACN concentration was then reduced to below 10%. Proteins and zelenectide pevedotin were digested overnight at 37 °C using a Trypsin/Lys-C mix at a protein-to-enzyme ratio of 1:20. Samples were centrifuged at 2000 g for 1 minute, and digestion was quenched by acidification with FA to a final concentration of 1%. Peptide solutions (200 ng in 1 μL) were injected into the LC-PRM system. The MMAE quantifier ion at *m/z* 686.5 was used for all spatial quantifications.

In the absence of a stable isotope-labeled internal standard for free MMAE, calibration curves were constructed using the pure zelenectide pevedotin reference standard. Quantitation of microdissected TC and TS regions was based on the diagnostic MMAE-derived fragment ion (*m/z* 686.5), representing the primary toxophore signature. Consequently, PRM peak areas were converted into estimated zelenectide pevedotin-equivalent amounts of payload on-column (fmol) using the external calibration curve for all spatial comparisons presented in Section 2.4. Method validation confirmed excellent linearity (*R*^2^ > 0.998) and high precision (*CV* ≤ 16% near the LOQ), establishing that this framework provides a robust and reproducible basis for relative micro-spatial pharmacokinetics.

Supplementary Table A1 reports both the raw PRM peak areas (AUC) and the corresponding estimated analyte amounts for all TC and TS regions across the four sampling timepoints.

### DIA-based spatial proteomic analysis for NECTIN-4 quantitation

Spatial proteomic analysis was performed on peptide digests generated from the remaining material obtained following laser capture microdissection (LCM) of tumour cell-rich (TC) and tumour stroma-rich (TS) regions. Samples corresponding to the same LCM-derived regions analysed by LC-PRM for spatial MMAE quantification were subjected to a complementary deep proteomics analysis based on data-independent data acquisition (DIA), enabling the relative quantitation of NECTIN-4 on tissue samples.

One microlitre (μL) of peptide digests were loaded on Evotips (EV2011, Evosep) as per the manufacturer’s protocol. Briefly, tips were rinsed with 20 µL 99.9 % acetonitrile 0.1 % formic acid by centrifugation, conditioned by soaking in 1-propanol, equilibrated with 20 µL Solvent A (0.1 % formic acid in water) by centrifugation, loaded with 1 µL of sample and 19 µL of solvent A, then centrifuged, washed with 20 µL solvent A by centrifugation, and wetted with 100 µL solvent A and a brief centrifugation step for 10 seconds. All centrifugation steps were 60 seconds at 800 g unless stated.

Peptides were analysed on an Evosep One LC system (EvoSep) coupled to a timsTOF Flex mass spectrometer (Bruker) using the 60 samples-per-day method and a 150 µm × 80 mm C18 column with 1.5 µm particles (Evosep, EV1109) and a 20 µm Captive Spray Emitter (Bruker). Buffer A was 0.1% formic acid in water and Buffer B was 0.1% formic acid in acetonitrile. Data were collected using diaPASEF with 1 MS frame and 8 diaPASEF frames per cycle with an accumulation and ramp time of 100 ms, for a total cycle time of 0.95 seconds. The diaPASEF frames were separated into 3 ion mobility windows, in total covering the 400–1000 *m/z* mass range with 25 *m/z*-wide windows between an ion mobility range of 0.65-1.45 Vs/cm^2^. The collision energy was ramped linearly over the ion mobility range, with 20 eV applied at 0.6 Vs/cm^2^ to 59 eV at 1.6 Vs/cm^2^.

Mass spectrometry raw files were analysed in DIA-NN version 2.3.1 using an in-silico spectral library generated by DIA-NN for precursor charge: 2-4 and precursor *m/z*: 400 - 1,000. Carbamidomethylation of cysteine was not selected and up to 2 missed cleavages were allowed. A Uniprot human + mouse FASTA file containing 20412 + 17246 sequences, with common contaminants added by DIA-NN was used as input to the spectral library prediction. Mass accuracy tolerances for MS1 and MS2 scans were set to 15 ppm, and scan window was set to 0 (automated assessment by DIA-NN), with the FDR threshold set to 1 %. The additional options “--pg-level 2” and “--species-genes” were used. Protein intensities from DIA-NN’s “pg_matrix” were used in further downstream analysis.

Protein abundance values for NECTIN-4 were extracted from the processed DIA dataset and compared with the corresponding LCM-LC-PRM measurements of MMAE obtained from matched tumour regions and post-dose timepoints. The DIA dataset was used exclusively to assess potential associations between local NECTIN-4 abundance and spatial MMAE distribution.

### Mass spectrometry data processing and analysis

Xcalibur software (version 4.7, Thermo Fisher Scientific) was used for instrument control and raw data acquisition. Raw files were processed and PRM peak areas (AUCs) for MMAE were extracted using Skyline (version 24.1, MacCoss Lab, University of Washington, Seattle, WA), configured for small-molecule targeted analysis [43], for both bulk tumour and LCM-derived tissue samples. Precursor and product ion information was imported to enable accurate transition selection and quantification. Statistical analyses and visualisations were performed using GraphPad Prism (version 10.4.2) and R package (v4.5.0). Chemical structures were generated using ChemDraw (version 22.2.0, PerkinElmer Informatics). Untargeted proteomic profiling of bulk tumour lysates was analysed using MaxQuant (v2.7.5.0), while the LCM-derived spatial proteomic data acquired by diaPASEF were analysed using DIA-NN (version 2.3.1). Heatmaps were generated using the pheatmap R package.

Statistical Analysis and Spatial Correlations: Bivariate correlation analyses between matched DIA-derived NECTIN-4 abundances and LCM-LC-PRM MMAE levels were performed. Pearson parametric correlation (r) and Spearman non-parametric rank correlation (*ρ*) coefficients were calculated. To prevent bias from missing value imputation, samples with non-detectable NECTIN-4 signal (ND) were excluded from bivariate correlation analyses, restricting the evaluated dataset to quantifiable ROIs (*n* = 17 out of 24 total micro-dissected regions (70.8%)). Two-tailed p-values and associated t-statistics were calculated using degrees of freedom d*f* = *n* - 2.

## ACKNOWLEDGEMENTS

We gratefully acknowledge Bicycle Therapeutics for their financial support during the conduct of this study. We also thank members of the Discovery Proteomics Facility and the Fischer & Kessler groups for critical and insightful comments on the manuscript.

## FUNDING

This study was funded by Bicycle Tx Limited. and supported by their research initiatives. The B.M.K. lab has been funded by the Chinese Academy of Medical Sciences (CAMS) Innovation Fund for Medical Sciences (CIFMS), China (grant number: 2024-I2M-2-001-1).

## AUTHOR CONTRIBUTIONS

M.K.I. and B.M.K designed the research. M.K.I. developed the methodology, prepared and analysed the samples, performed all experiments, and operated the LC-MS/MS system, including targeted LC-PRM, for data acquisition and analysis. S.D. and T.H. provided expert assistance with the LCM microdissection. S.H. provided expert guidance with the PRM development protocol. M.K.I. drafted the initial manuscript, and M.K.I. and B.M.K. jointly revised and finalised the text. M.K.I., S.D., T.H., S.H., M.R., F.H., P.J., M.S., R.F. and B.M.K. all provided critical comments and contributed to the editing of the manuscript.

## DECLARATION OF COMPETING INTERESTS

M.R., F.H., P.J. and M.S. are employed by Bicycle Therapeutics. The remaining authors declare no competing interests.

## AI USE DECLARATION

We have used ChatGPT (EDU, University of Oxford; GPT-5.6 Sol) for minor text drafting, content summarisation, and text editing.

## DATA AVAILABILITY

The data generated and evaluated in this manuscript are included in the published article and its Supplementary Information. Raw and meta MS data will be uploaded and deposited onto PRIDE.

## Reviewer access details

Log in to the PRIDE website (https://www.ebi.ac.uk/pride/) using the following details:

DDA and PRM data for MMAE quantitation: Will be uploaded to PRIDE.

DIA-MS data for NECTIN-4 quantitation: Project accession: PXD083605; Token: MNjBFL33l9Fq

## Notes

### Competing Interest Statement

The authors have declared no competing interest.

