## Supplementary Information for "Spatial Tumour Tissue Distribution Dynamics of a Bicycle Peptide-MMAE Toxin Conjugate Profiled by Targeted Mass Spectrometry"

**Supplementary Figures, Tables and Figure Legends**

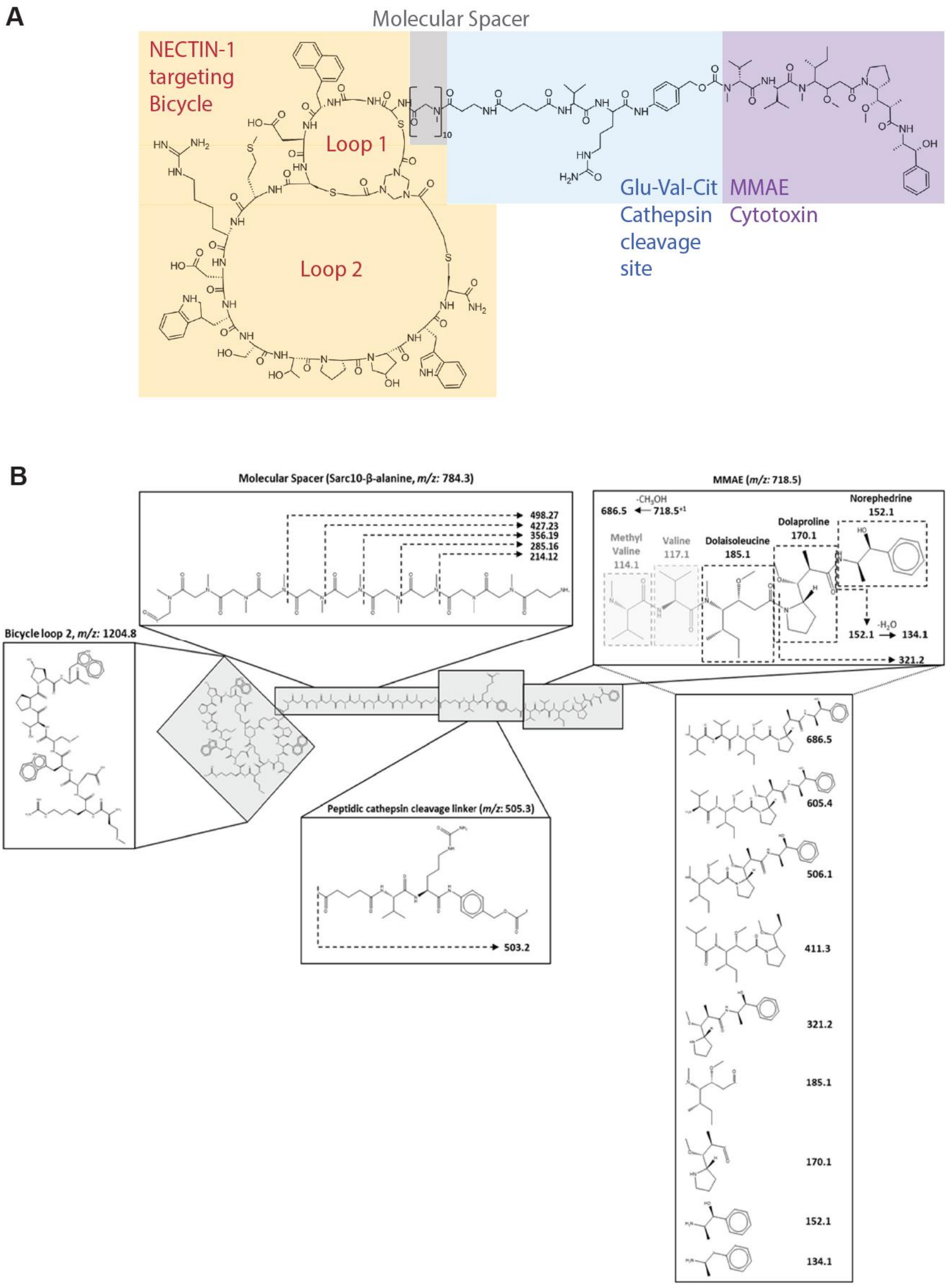


**Figure S1. (A) Chemical structure, and (B) proposed fragmentation profile of zelenectide pevedotin.** Fragment ions observed by MS/MS are proposed to originate from specific regions of the molecule (namely the bicycle structure, spacer, MMAE moiety, and linker) based on manual structural annotation using ChemDraw and comparison with published fragmentation patterns, with the respective fragment ions assigned to their origins as described in the main text. Cleavage sites are marked according to characteristic features, and each fragment ion is mapped accordingly. This schematic provides a qualitative interpretation of zelenectide pevedotin fragmentation to support analysis of mass spectral data.

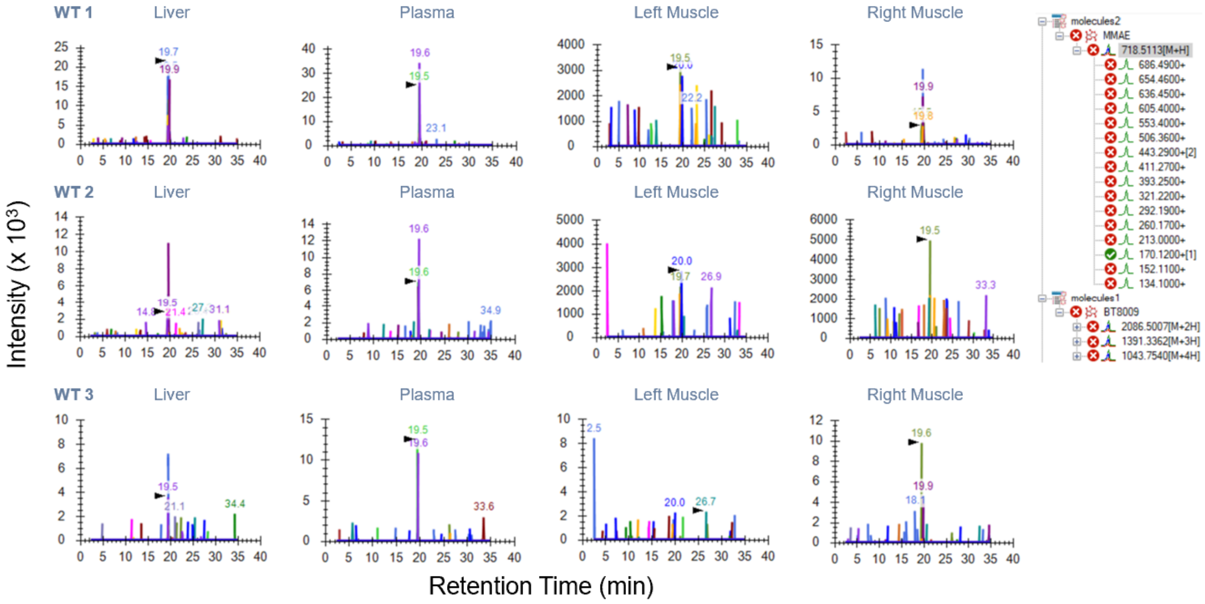


**Figure S2. Skyline-extracted chromatograms from tissues of wild-type (WT) control mice.** No zelenectide pevedotin-related fragment ions were detected in the analysed samples, supporting the specificity of the LC-PRM-MS/MS method.

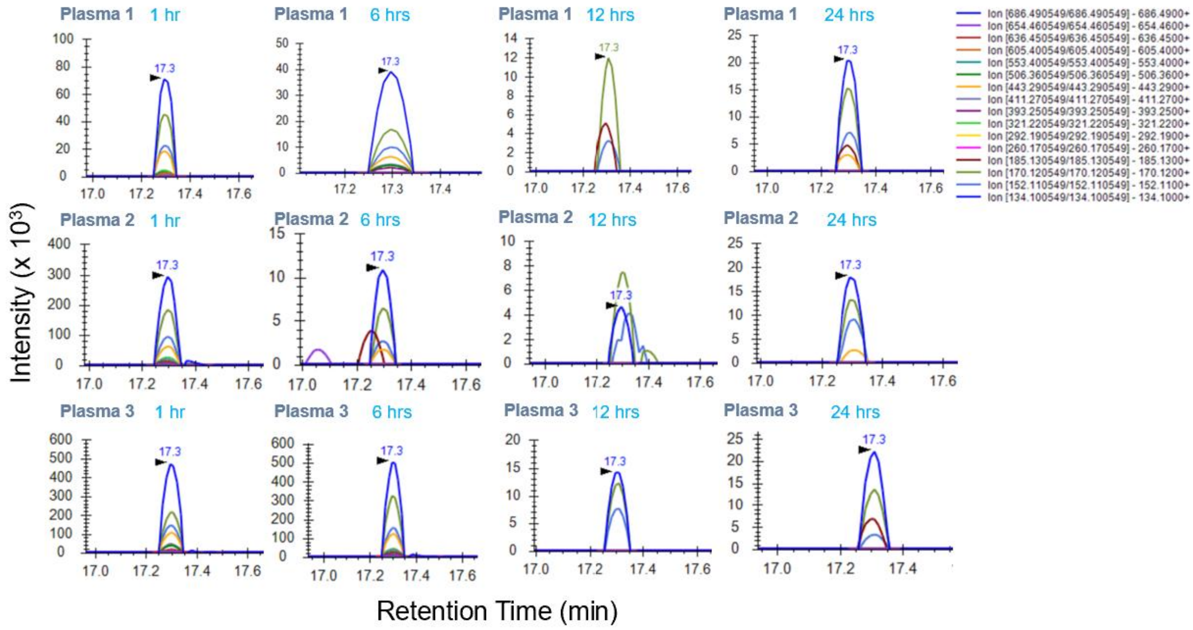


**Figure S3. LC-PRM-MS/MS chromatograms of plasma samples collected at 1, 6, 12, and 24 h post-administration of zelenectide pevedotin.** Each chromatogram shows the fragmentation profile of monomethyl auristatin E (MMAE) in plasma, as analysed using Skyline. The fragment ion at *m/z* 686.5 exhibits the highest signal intensity among the monitored ions across all displayed time points. Signal intensity reflects dynamic systemic exposure over time, supporting effective drug circulation and clearance.

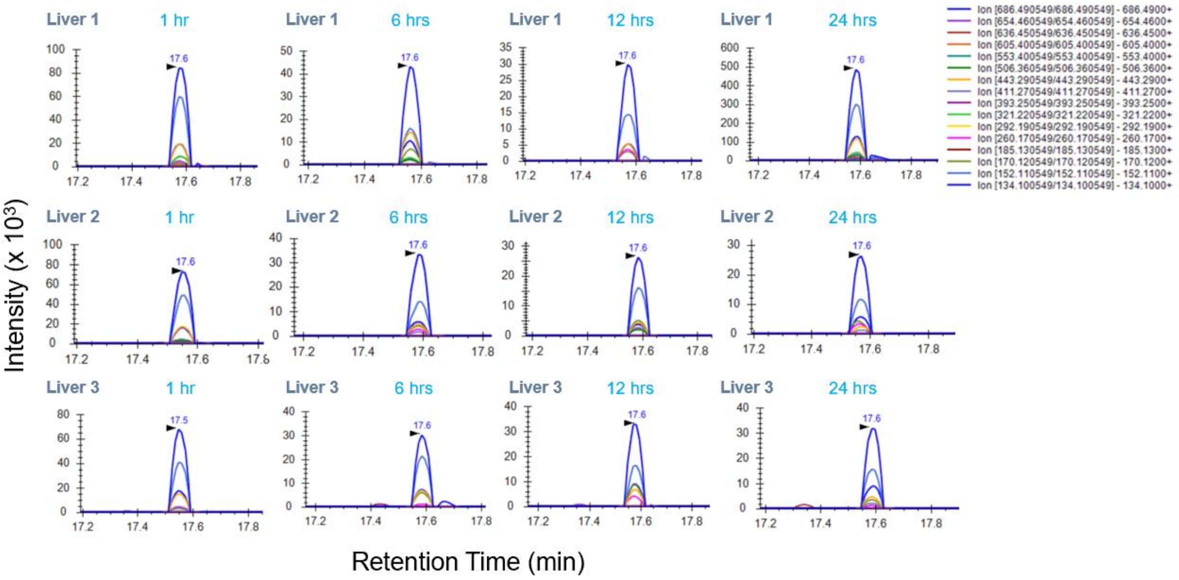


**Figure S4. LC-PRM-MS/MS chromatograms of liver samples collected at 1, 6, 12, and 24 h post-administration of zelenectide pevedotin.** The chromatograms show the fragmentation profile of monomethyl auristatin E (MMAE) in liver tissue across the indicated time points. The fragment ion at *m/z* 686.5 is observed across the displayed time points.

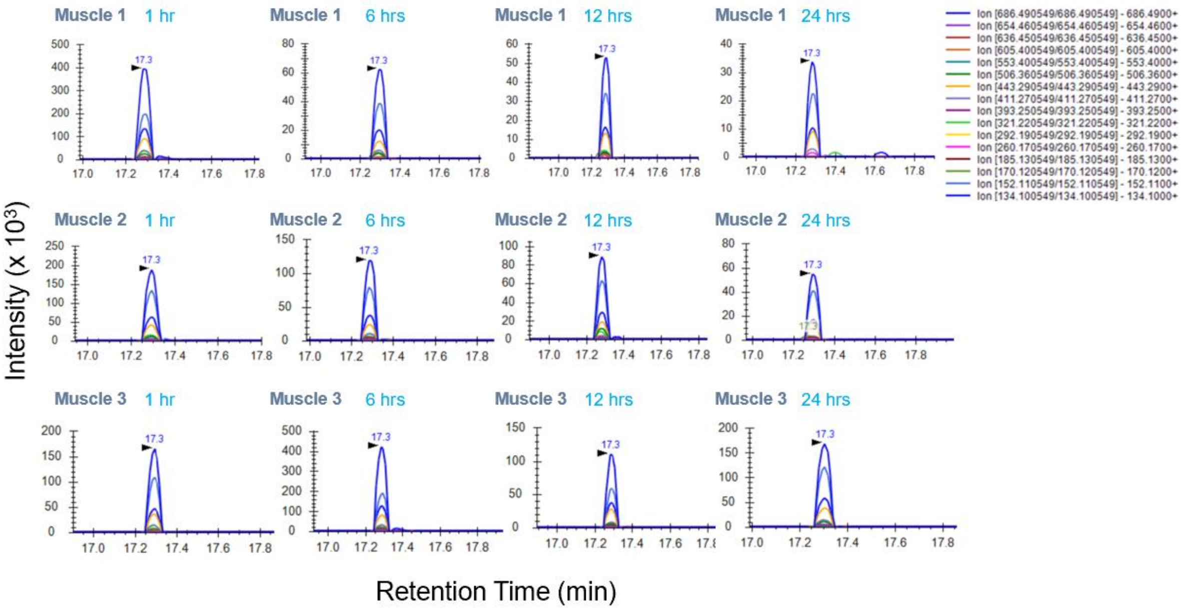


**Figure S5. LC-PRM-MS/MS chromatograms of muscle samples collected at 1, 6, 12, and 24 h post-administration of zelenectide pevedotin.** The chromatograms show the fragmentation profile of monomethyl auristatin E (MMAE) in muscle tissue across the indicated time points. The consistent intensity and retention of the key fragment ion at *m/z* 686.5 validate the PRM method's reliability in low-abundance matrices. The observed profile suggests limited accumulation of MMAE in this non-target tissue.

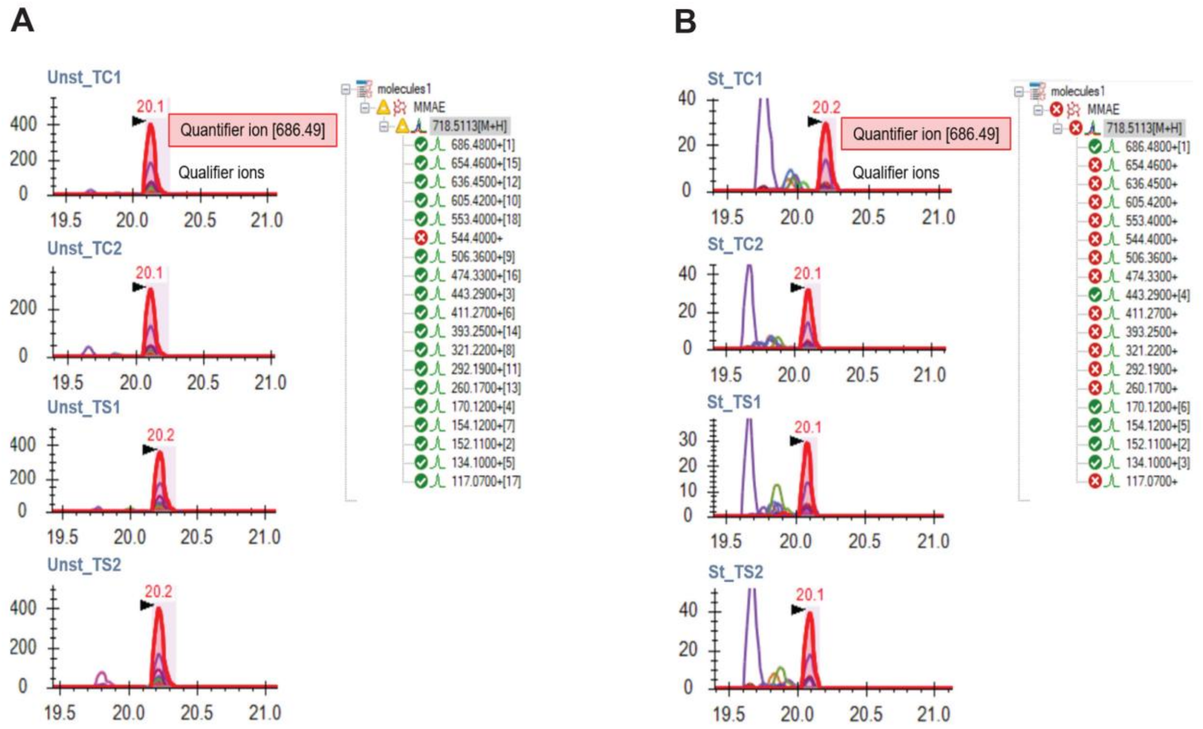

**Figure S6. Comparison of MMAE detectability between haematoxylin and eosin (H&E)-stained and unstained tumour cryosections analysed by LCM-PRM-MS/MS.** Tumour tissue collected 12 h post-dose-the time point with the highest MMAE abundance in bulk PRM-MS/MS-was used to generate unstained (panel A) and H&E-stained (panel B) 5 µm cryosections. Four regions of interest (ROIs) were selected per condition: two tumour cell-rich regions (TC1, TC2) and two stroma-rich regions (TS1, TS2), isolated by LCM and processed identically for PRM-MS/MS analysis. Unstained sections consistently produced higher MMAE quantifier-ion signal intensities and allowed robust detection of all MMAE qualifier ions, whereas H&E-stained sections showed reduced signal and diminished sensitivity. These results indicate that histological staining can negatively impact MMAE detectability in spatially resolved LC-MS/MS analyses. ROI labels indicate staining status (Unst = unstained; St = H&E-stained) and tissue category (TC = tumour cell-rich; TS = tumour stroma-rich).

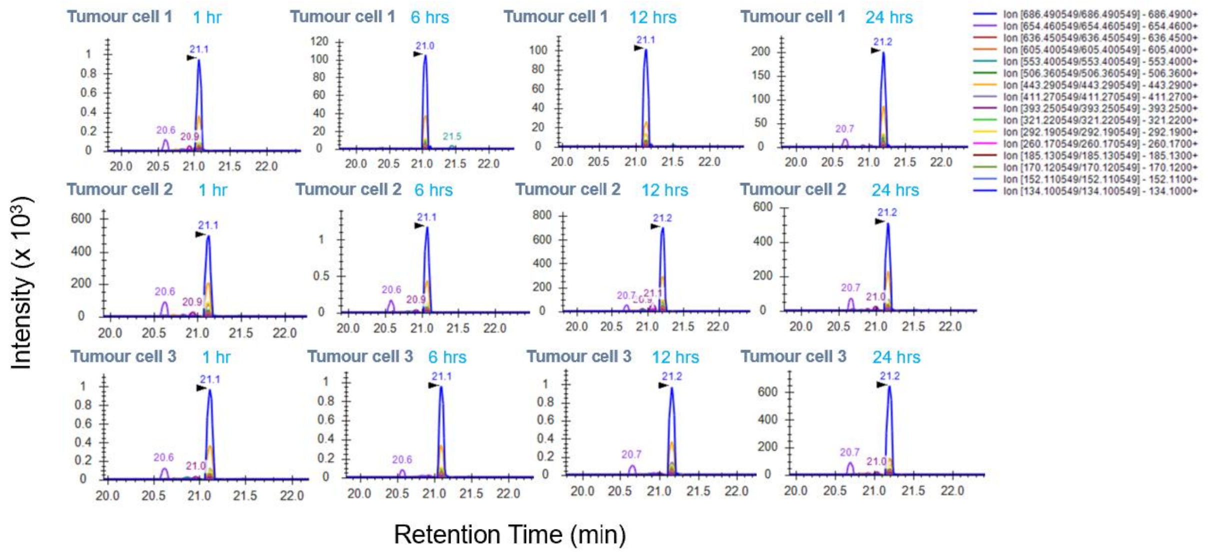

**Figure S7 LCM-LC-PRM-MS/MS chromatograms of tumour cell samples collected 1, 6, 12, and 24 h post-administration of zelenectide pevedotin.** Each chromatogram shows the fragmentation profile of monomethyl auristatin E (MMAE) within tumour cell regions. The fragment ion at *m/z* 686.5 exhibits the highest signal intensity among the monitored ions across all displayed time points, highlighting the method's reproducibility and sensitivity within tumour subregion matrices.

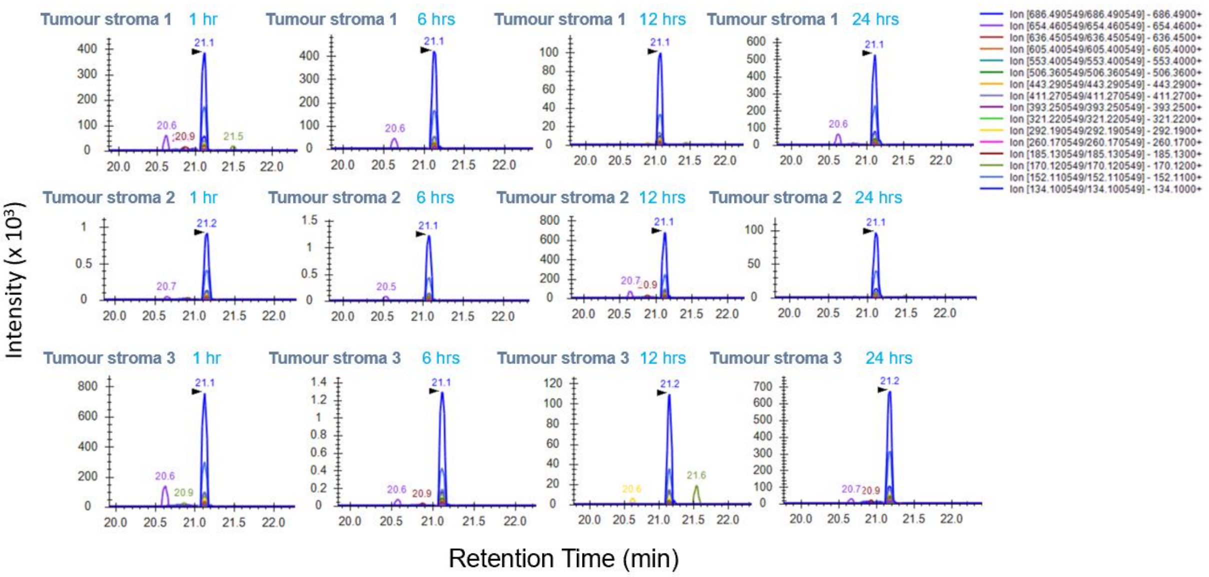


**Figure S8 LCM-LC-PRM-MS/MS chromatograms of tumour stroma samples collected at 1, 6, 12, and 24 h post-administration of zelenectide pevedotin (MMAE).** Each chromatogram shows the fragmentation profile of monomethyl auristatin E (MMAE) within tumour stroma regions. The fragment ion at *m/z* 686.5 across all time points supports the method's sensitivity and reproducibility in complex tumour microenvironment matrices.

### 1 Supplementary Information for Materials and Methods Section

**Table S1.** Treatment groups and corresponding sample sizes. Group 1 consisted of wild-type (WT) mice. Groups 2 to 5 received zelenectide pevedotin, with samples collected at **1, 6, 12, and 24 h** post-administration, respectively.

| | | | | Tissue Type | | | |
|---|---|---|---|---|---|---|---|
| **Group** | **Treatment** | **Sampling (after dose)** | **Dosage** | **Plasma** | **Liver** | **Tumour** | **Gastrocnemius (Skeletal Muscle)** |
| 1 | wt | - | --- | 3 | 3 | - | 3 |
| 2 | zelenectide | 1h | 3mpk | 3 | 3 | 3 | 3 |
| 3 | zelenectide | 6h | 3mpk | 3 | 3 | 3 | 3 |
| 4 | zelenectide | 12h | 3mpk | 3 | 3 | 3 | 3 |
| 5 | zelenectide | 24h | 3mpk | 3 | 3 | 3 | 3 |
| | | | **Total** | **15** | **15** | **12** | **15** |

**Table S2**. The inclusion list for zelenectide pevedotin MS2 data generation using ions corresponding to MS1 data.

| **Mass [*m/z*]** | **CS [*z*]** | **Polarity** | **(N)CE** |
|---|---|---|---|
| 2087.5 | 2 | Positive | 44 |
| 1392.3 | 3 | Positive | 44 |
| 1044.7 | 4 | Positive | 44 |
| 718.5 | 1 | Positive | 28 |

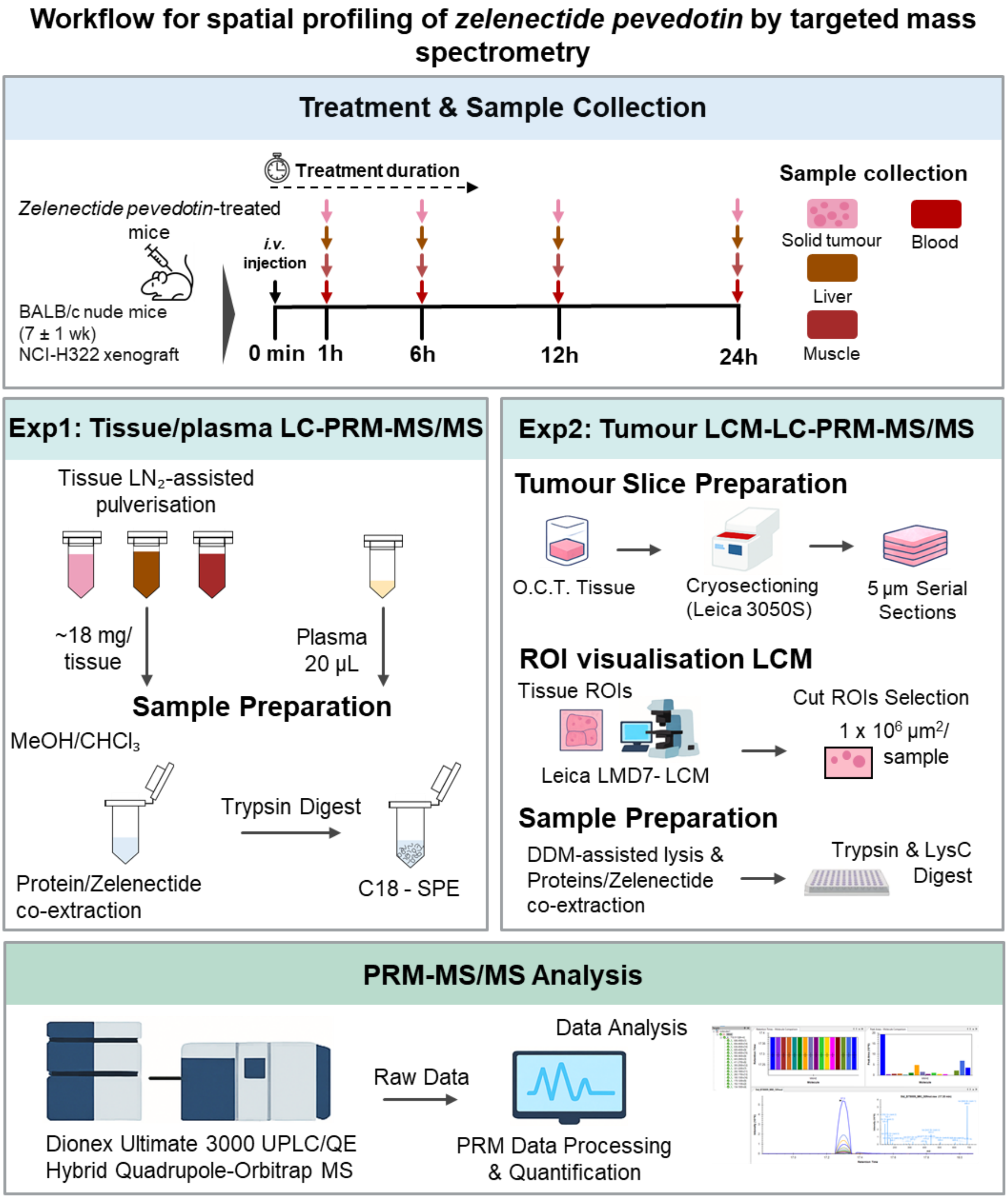


**Figure S9. Integrated workflow for spatial profiling of zelenectide pevedotin using targeted mass spectrometry.** Schematic overview of the complete analytical pipeline. BALB/c nude mice (7 ± 1 wk) were treated with zelenectide pevedotin and tissues (solid tumour, liver, muscle, plasma) collected at 6 time points. **Experiment 1:** ~18 mg of each tissue were homogenised in liquid nitrogen (plasma: 20 µL directly), followed by MeOH/$CHCl_3$ co-extraction, trypsin digestion, and solid-phase (SPE) extraction using C18 column. **Experiment 2:** OCT-embedded solid tumours were cryosectioned (5 µm), regions of interest (ROIs) (~$1 \times 10^6$ µm$^2$) visualised and captured by laser capture microdissection (LCM), lysed using DDM-assisted digestion, followed by trypsin/LysC digestion. Samples from both experiments were analysed by targeted PRM-MS/MS for quantitation of zelenectide pevedotin.
